# Cell-cell communication analysis demonstrates early-stage common pathways linking ageing, Alzheimer’s disease, and Type 2 diabetes-related brain dysfunction

**DOI:** 10.1101/2025.10.24.684393

**Authors:** Boyong Wei, Alan Stitt, Karis Little

## Abstract

Ageing promotes to the development of age-related diseases and cognitive decline. In this study, we integrated and analysed four single-cell RNA sequencing (scRNA-seq) datasets encompassing Alzheimer’s disease, type 2 diabetes, and ageing in mouse brain tissue to identify early pathological factors that may drive normal ageing toward disease through alterations in cell–cell communication (CCC). Building on our previously established CCC change modelling framework, we found that both Alzheimer’s disease and ageing were characterized by a loss of intercellular communication, whereas type 2 diabetes exhibited an overall gain of new communication pathways. Notably, vascular communication changes were more prominent in age-related diseases than in normal ageing. Furthermore, we identified a series of CCC molecules that play key roles in brain ageing and disease through pseudo covariance and conflicting resolving CCC algorithms. Among them, the LRP1 receptor on astrocytes emerged as a central CCC hub implicated in both ageing and disease pathology.

## Introduction

Ageing is a highly complex biological process encompassing various hallmarks, including genomic instability, telomere attrition, epigenetic alterations, and altered CCC (López-Otín et al., 2013). Despite the well-established link between age-related pathogenic processes and age-related diseases, identifying the key mechanisms driving this relationship remains challenging. Since ageing is a time-dependent process, it is reasonable to infer that certain molecular and cellular changes occurring during ageing may represent early-stage drivers or even cause age-related diseases. If such changes exist, they could serve as potential therapeutic targets for preventive strategies. It has been shown that cognitive decline associated with ageing, T2D, and AD is associated with neurovascular pathology, especially linked to progressive dysfunction of the neurovascular unit (NVU) (Barloese et al., 2022; S. Liu et al., 2023).

It is now widely appreciated that there is strong connectivity between seemingly disparate tissues. This can manifest in chronic disease where multiple organs are affected resulting in multi-morbidity. In particular, age-associated conditions such as Type 2 diabetes (T2D), Alzheimer’s disease (AD), and cardiovascular disease are often linked through shared pathogenesis and co-morbidities (J. Guo et al., 2022). Meta-analyses using data from a UK biobank has demonstrated that T2D can accelerate brain ageing (Dove et al., 2024) and cognitive decline (Antal et al., 2022)(Y. Liu et al., 2024). Numerous epidemiological studies have demonstrated that T2D patients have a significantly higher risk of developing Alzheimer’s disease (AD) in comparison with age-matched non-diabetic subjects (Cummings et al., 2022; Sridhar et al., 2015). The number of cases of dementia associated with T2D is expected to increase because of the diabetes pandemic and the concomitant rise in ageing populations worldwide.

The NVU is crucial for integrity of the CNS. In the brain, the normal function of the NVU relies heavily on complex and dynamic CCC between component cells to maintain tissue homeostasis across the lifespan (S. Guo & Lo, 2008). There has been focused interest on the critical role of astrocyte interactions with the vasculature, neurons and immune cells (Huang et al., 2019; Manu et al., 2023; Mishra & Kumar, 2025; Rustenhoven et al., 2016; Sweeney et al., 2016). Normal CCC within the NVU can become disturbed in pathological settings. This altered intracellular crosstalk has been implicated in neurodegenerative conditions ranging from AD disease, Parkinson’s disease, multiple sclerosis and diabetic retinal disease (Bury et al., 2021; Chaudhuri et al., 2025).

In this study, a number of datasets were analysed in relation to CCC in order to identify potential driving factors behind age-related diseases. An unbiased, dual analysis of single-cell RNA sequencing (scRNA-seq) datasets was conducted from both ageing and age-related disease studies (2 Alzheimer’s diseases datasets, one type 2 diabetic dataset, one ageing dataset), with a focus on defining CCC alterations. We utilised our previously established CCC modelling method (REF) and generated the significant CCC categorical changes in each dataset. Based on those categorical changes, we developed a pseudo covariance algorithm to converge the age-related diseases CCC changes.

We also introduced a novel concept on ‘conflict’ of CCC change (see *overlapping changes* and *Prevention action inference* in methods section). This notion arose from that the nature of CCC is between entities (source cell type to give the ligand as the sender, the target cell type to have the receptor as receiver). Thus, the same molecule on the same cell type can have different directional changes in different sender and receiver CCC pairs. Our ageing mapping algorithm that incorporated this feature, used ageing changes as negative reference to infer the actions to correct the trend from normal ageing going towards pathological conditions.

The REF in this preprint refers to the CCC modelling paper we previously established but is currently under revision peer review.

## Methods

The first methodological step was modelling the convergence between the impact of T2D and AD on the brain, and then mapping these findings onto ageing-related changes (Fig 1). While the mapping process was complex, the core idea was straightforward: assuming that most ageing-associated changes are detrimental, it considered changes occurring in the opposite direction to be beneficial or indicative of proper CCC function.

**Figure 1.**
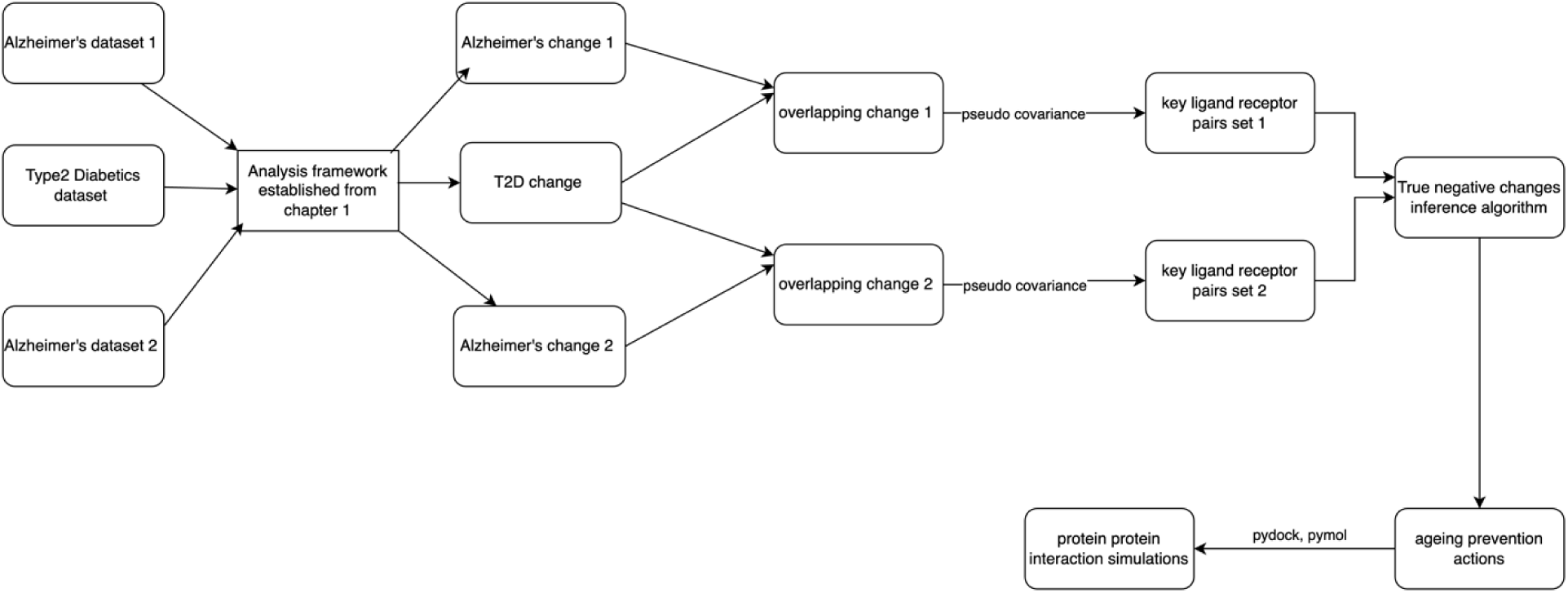
Workflow of this study. Three scRNA dataset integrative analysis was conducted first, which was later mapped to ageing dataset to generate ageing prevention actions.

### Dataset collection info

The ageing dataset was obtained from (Almanzar et al., 2020), while the h5ad data was downloaded from the CellxGene platform (https://cellxgene.cziscience.com/collections/0b9d8a04-bb9d-44da-aa27-705bb65b54eb). This dataset, titled "Brain non-myeloid cells – A single-cell transcriptomic atlas characterizes ageing tissues in the mouse", provides a comprehensive single-cell transcriptomic analysis of ageing mouse brain. The T2D-related dataset was sourced from (Little et al., 2023) and is publicly available under GEO accession GSE217665. For AD disease models, two datasets were utilized: AD_6 was obtained from (Park et al., 2023) under GEO accession GSE224398, where only the 6-month data subset was included in the analysis. AD_7 was derived from (Fatmi et al., 2024) GSE227157, with only data from the AD model and control groups being used. Each dataset was carefully selected to ensure relevance and consistency across studies, facilitating a robust comparative analysis of ageing, T2D, and AD disease at the single-cell level.

### Gene translation

For the ageing dataset, the raw count matrix from h5ad object downloaded was extracted and then turned into a scanpy(1.9.6) object. After that, the object was used as input for mousipy (0.1.3) translate function. The translation result was ‘Found direct orthologs for 16154 genes. Found multiple orthologs for 713 genes. Found no orthologs for 4202 genes. Found no index in biomart for 0 genes.’

For T2D dataset, the scanpy(1.9.6) object was directly created from downloaded 10x mtx format data using read_10x_mtx function. After that, the object was used as input for mousipy (0.1.3) translate function. The translation results in both control and T2D conditions were the same ‘Found direct orthologs for 16787 genes. Found multiple orthologs for 781 genes. Found no orthologs for 8234 genes. Found no index in biomart for 2196 genes.’

For AD_6 dataset, the scanpy(1.9.6) object was created through read_10x_h5 as the downloaded data format was in h5. After that, the object was used as input for mousipy (0.1.3) translate function. The translation result in control was ‘Found direct orthologs for 17483 genes. Found multiple orthologs for 976 genes. Found no orthologs for 13137 genes. Found no index in biomart for 689 genes.’ The translation result in AD was ‘Found direct orthologs for 16801 genes. Found multiple orthologs for 697 genes. Found no orthologs for 5216 genes. Found no index in biomart for 1416 genes.’

For AD_7 dataset, the scanpy(1.9.6) object was directly created from downloaded 10x mtx format data using read_10x_mtx function. After that, the object was used as input for mousipy (0.1.3) translate function. The translation results in both control and T2D conditions were the same ‘Found direct orthologs for 17483 genes. Found multiple orthologs for 976 genes. Found no orthologs for 13137 genes. Found no index in biomart for 689 genes.’

### scRNA pipeline

All datasets used the same pipeline between translated scanpy(1.9.6) object and clustered object. The first step was var_names_make_unique function to make sure there were no duplicates in the gene dimension. Gene dimensions that had fewer than 3 cells and cells that had less than 200 genes were all removed. Next, the cells that had more than 10 percent mitochondrial gene counts were removed. After that, the counts per cell was normalized to 10000 and then log1ped using sc.pp.normalize_total and sc.pp.log1p functions. The highly variable genes were calculated by sc.pp.highly_variable_genes with min_mean=0.0125, max_mean=3, min_disp=0.5. Then the raw object was saved and the object was filtered to only highly variable gene dimensions. The sc.pp.regress_out was applied to regress out of mitochondrial percentage. The unit variance was scaled using sc.pp.scale with max_value=10. Finally, PCA was conducted using sc.tl.pca function with svd_solver=’arpack’, and a UMAP was generated with n_neighbors=10, n_pcs=30 by sc.pp.neighbors function. The data was clustered by sc.tl.leiden function with resolution = 0.2.

After clustering, in the ageing dataset, the clusters which contained lower than 150 cells were removed. In the T2D dataset, the clusters that had lower than 200 cells in both control and T2D conditions were discarded. In the AD_6 dataset, the clusters lower than 200 cells in control were discarded and the clusters lower than 149 cells in AD condition were eliminated. In the AD_7 dataset, the clusters that had lower than 200 cells in both control and AD conditions were discarded. The thresholds are determined manually by manual decision on the cluster shape.

The exact annotation proofs are shown in (Sup Table S2).

### CCC inference

The cell to cell communication activities from annotated scanpy objects were inferred from LIANA (0.1.9). This version of LIANA still gave two p values (cellphonedb and cellchat (Efremova et al., 2020; Jin et al., 2021)) which later versions discarded. The exact function and parameters used were rank_aggregate.__call__, with groupby=’cell_type’, resource_name=’consensus’, expr_prop= 0.1, min_cells = 5, base = 2.718281828459045, aggregate_method=’rra’, return_all_lrs = False, consensus_opts=None, use_raw = True, layer = None, de_method=’t-test’, verbose = False, n_perms = 1000, seed = 1337, resource = None, inplace=False.

After the function returned a result, it was filtered by double p values through 0.05 threshold. Only the filtered result was used for following analysis.

### CCC change analysis

The CCC change analysis derived from CCC inference was the same as described in our previously published paper (REF). Only the shortlisted changes were proceeded to the further steps.

### Commonly changed LR pairs between T2D and AD

The term "commonly changed ligand-receptor (LR) pairs" refers to LR pairs that exhibit alterations in both T2D and AD conditions. This means that if a particular LR pair, for example APP–LRP1, is identified as a commonly changed pair between the T2D and AD6 datasets, it indicates that APP-LRP1 underwent some form of communication change in both conditions. However, this designation does not imply that the direction or nature of change (e.g., upregulation or downregulation) was necessarily the same across both disease states. Instead, it highlights molecular interactions that were affected in both pathological conditions. The primary objective of this study was to uncover underlying factors contributing to age-related diseases by identifying molecular disruptions common to multiple conditions. Therefore, changes that were specific to only one pathological condition (either T2D or AD) were not the primary focus, as they do not provide insights into shared disease mechanisms. Instead, emphasis was placed on identifying molecular interactions that may play a central role in ageing-associated diseases by being altered in multiple disease contexts.

### Pseudo covariance algorithm

The pseudo-covariance algorithm was specifically designed to quantify the degree of agreement or opposition between ligand-receptor (LR) pair changes across overlapping communication types. The primary unit of analysis was the ligand-receptor pair, while the variable was the type of communication, each associated with a categorical change: gain, loss, or consensus. The algorithm assigned numerical values to different interaction patterns to capture the directionality and coherence of changes. If two communication types both exhibited consensus changes, their contribution to the pseudo-covariance was 0.5, reflecting a neutral effect. When a single consensus change was paired with either a gain or loss, the assigned product was 0.25, indicating partial agreement. However, in cases where no consensus change was present, the interaction was categorized based on directional agreement. If both conditions (e.g., T2D and AD) showed the same type of change, either two gains or two losses, the assigned product was 1, reflecting complete alignment. Conversely, if one condition exhibited a gain while the other showed a loss, the product was -1, indicating a strong opposing trend. To compute the final pseudo-covariance value for a given ligand-receptor pair, the algorithm summed the assigned products across all overlapping communication types. This cumulative score served as a measure of the degree of alignment or discordance between disease-related changes, providing a structured approach to quantify the interactions between overlapping ligand-receptor communication patterns. The resulting values helped in determining whether a particular ligand-receptor pair exhibited coordinated or conflicting changes across the pathological conditions under study.

### Overlapping changes

Building upon the concept of commonly changed ligand-receptor (LR) pairs, overlapping changes refer to LR pairs that not only exhibit alterations in both T2D and AD conditions but also share the same type of communication. This means that in both pathological states, the interaction occurs between the same source cell type (expressing the ligand) and the same target cell type (expressing the receptor). Overlapping changes are characterized by three key elements:

1. Type of Communication – Identifying which source cell type expresses the ligand and which target cell type expresses the corresponding receptor.
2. Ligand-Receptor Pair – The specific molecular interaction involved in the communication.
3. Change Direction – The nature of the alteration observed across both pathological conditions, classified into one of three categories: Consensus Change – The LR pair exhibits a moderate trend in the condition (e.g., upregulated or downregulated). Gain – The interaction is newly established in the condition, indicating a previously absent communication that emerges in the disease state. Loss – The interaction is disrupted in the condition, suggesting a communication pathway that is diminished or eliminated in the disease state.

By focusing on overlapping changes, the study aims to identify shared disruptions in cellular communication that may contribute to age-related disease progression. This approach helps to distinguish molecular interactions that could serve as potential therapeutic targets or biomarkers for age-related pathologies.

### Prevention action inference

Inferring prevention actions in the context of CCC changes is a highly complex process, requiring careful consideration of multiple interacting factors to determine the correct direction of change and resolve potential conflicts. Several challenges arise when attempting to infer the optimal direction for CCC modifications across different pathological conditions.

Key Challenges in Prevention Action Inference included roughly three factors. The first was involvement of multiple conditions and the role of ageing as an anchor. The inference process involves three conditions: ageing, T2D, and AD disease. To establish a reference point, the direction of change in ageing is used as an anchor, under the assumption that the opposite of the ageing-related change represents a beneficial or preventive direction. If an interaction gains with ageing, the presumed good direction would be a loss of that interaction. Conversely, if an interaction declines with ageing, maintaining or restoring it may be beneficial. This assumption, however, introduces complexity when interactions exhibit inconsistent changes across different conditions. Secondly, same Ligand-Receptor Pair with different types of communication, one ligand-receptor (LR) pair can be involved in multiple types of communication, where it can have different change directions. In some cases, the ageing-based inferred direction for one type of communication may contradict the direction inferred for another type. This creates a challenge in determining whether a uniform prevention action can be applied across all instances of that LR pair or if context-specific adjustments are required. Third, interactions with Overlapping and Non-Overlapping Changes. Some LR pairs show overlapping changes between T2D and AD, meaning they share both the type of communication and the direction of change. However, ageing-related changes in an LR pair can also involve additional, non-overlapping interactions, where the same LR pair is altered in different types of communication that are not shared between T2D and AD. The challenge lies in deciding whether and how to incorporate these non-overlapping changes into the inference process, especially if they suggest opposing trends.

### Resolving Conflicts in Prevention Action Inference: A Step-by-Step Framework

To establish a robust prevention action inference framework and address conflicts systematically, a structured multi-step approach was implemented. This process ensures that changes in cell-cell communication (CCC) are correctly interpreted, and conflicting inferences are resolved before identifying final prevention strategies. Step 1: Pair-Based Processing of Overlapping Changes. All overlapping changes were processed in a ligand-receptor pair-based manner, meaning that all occurrences of the same ligand-receptor pair across different types of communication (i.e., different source and target cell types) were analysed together. Consensus changes in overlapping conditions were temporarily removed since they do not contribute to reversing pathological alterations or create conflicts. After filtering out consensus changes, five possible scenarios remained: Only one gain, Only one loss, Two losses, Two gains, One gain and one loss. The first four scenarios (one or two changes in the same direction) were straightforward, and the corresponding change was directly recorded as relevant. The one gain and one loss scenario was excluded from further consideration, as the contradictory trends suggest that the ligand-receptor interaction in that particular communication type is unlikely to be a primary driver of pathology. Step 2: Resolving Conflicts in Ageing Changes. After addressing overlapping changes in disease conditions, the next step was to process ageing-related changes, which were also handled in a ligand-receptor pair-based manner. Two ageing stages were considered: 3 to 18 months, 18 to 24 months. Similar to pathological conditions, conflicts between ageing stages were resolved: Consensus changes were filtered out. One gain and one loss scenario was removed, as the contradiction suggests an inconsistent pattern. The remaining ageing changes were then compared against their counterparts in pathological conditions to determine their relevance in prevention action inference. Step 3: Matching Overlapping Changes with Ageing Changes. Each overlapping change was checked for a corresponding change in ageing. If no counterpart was found in ageing, the overlapping change was by default classified as "bad" since it only appeared in disease conditions and was absent in normal ageing. The reverse direction of this change was recorded as a potential prevention action candidate. If the overlapping change had a counterpart in ageing with the same direction, the reverse direction of the overlapping change was still recorded as a prevention action candidate, since the alteration occurred in both ageing and disease conditions. If the overlapping change had a counterpart in ageing with the opposite direction, the overlapping change was deemed "good" or "benign", meaning no further action was necessary, and this interaction was discarded from further prevention inference. Step 4: Incorporating Non-Overlapping Ageing Changes. Although the primary focus was on overlapping changes, non-overlapping ageing-related changes (i.e., changes that occurred in ageing on the same ligand receptor pair but were not found in the same type of communication changes of disease conditions) were still considered. The reverse direction of these ageing changes was recorded as additional potential prevention action candidates since they may indicate beneficial modifications independent of pathological conditions. Step 5: Final Conflict Resolution for Prevention Actions. Once all potential prevention action candidates were gathered, a final conflict resolution step was performed. Candidates that conflicted with others were removed to ensure that the final set of net prevention actions contained only internally consistent strategies.

Example Illustration: Relationship Between Prevention Action Candidates and Ageing Change Direction. Consider the following scenario. Ageing Change: At the x,y coordinate [(endothelial *→* astrocyte), (PKM, CD44)], the interaction exhibits a loss direction after conflict resolution. The same x,y coordinate in ageing also shows a loss direction after resolving conflicts. Inference: the reverse direction of the ageing change is gain. Since the overlapping change in the disease conditions was a loss, the recommended prevention action is to increase PKM expression in endothelial cells and increase CD44 expression in astrocytes. This candidate consists of two components: Upregulating the ligand (PKM) in the source cell (endothelial cells), Upregulating the receptor (CD44) in the target cell (astrocytes). Final Conflict Resolution at the Single Action Level Instead of resolving conflicts at the ligand-receptor pair level, the final step operates at the single-action level. If any individual ligand or receptor action contradicts another inferred action, it is removed to maintain a consistent prevention strategy.

### Resveratrol Dataset Information

The study “Resveratrol preconditioning induces genomic and metabolic adaptations within the long-term window of cerebral ischemic tolerance leading to bioenergetic efficiency” (Khoury et al., 2018) investigated the effects of resveratrol preconditioning on cerebral ischemic tolerance. This dataset includes bulk RNA sequencing experiments conducted on 10-week-old mice, with resveratrol treatment (10mg/kg) administered from 8 weeks of age. The study aimed to explore genomic and metabolic adaptations that enhance bioenergetic efficiency in response to ischemic stress.

### Protein dock simulation

The structure of LRP1 used in this study was obtained from the RCSB Protein Data Bank (RCSB PDB) under the PDB ID: 2FYJ (Jensen et al., 2006). This structure was determined using nuclear magnetic resonance (NMR) and corresponds specifically to the CR56 domain of LRP1. The CR56 domain is particularly significant because it has been identified as containing key binding determinants for numerous LRP1 ligands, making it an ideal candidate for docking simulations aimed at inferring ligand-receptor interactions. Unlike LRP1, a convenient and structurally similar domain for APP (Amyloid Precursor Protein) was not available in the PDB database. As a substitute, the transmembrane domain of APP, with PDB ID: 2LLM, was used (Nadezhdin et al., 2011). This structure, also determined by NMR, provides valuable insight into the membrane-spanning region of APP, which plays a crucial role in its cleavage and processing, leading to the production of amyloid-beta (AB). While the extracellular domain of APP would have been ideal for ligand-receptor docking studies, its unavailability necessitated the use of the transmembrane domain as an alternative. On the other hand, numerous structures exist for amyloid-beta (AB) peptides, given their central role in AD disease pathology. Among these, the amyloid β-peptide (1–42) structure, available under PDB ID: 1IYT, was selected (Crescenzi et al., 2002). Like the other structures, 1IYT was determined through NMR spectroscopy and provides a high-resolution model of the AB (1–42) peptide, which is known for its propensity to aggregate and form amyloid plaques. Although from the scRNA CCC inference, C1QB with LRP1 was shortlisted, there is only c1q structure data in PDB dataset not specific to C1QB. Thus, the structure used to represent C1QB could be also representative for other protein that contained c1q. ‘5HKJ’ structure was chose as the paper indicated it senses a wide variety of ligands (Moreau et al., 2016). Saposin B structure which represented PSAP gene was ‘1N69’ (Ahn et al., 2002). Both ‘5HKJ’ and ‘1N69’ were measured by X-RAY DIFFRACTION. Together, these structural selections enabled a comprehensive molecular docking and binding interaction analysis between LRP1, APP, and amyloid-beta, facilitating insights into their potential roles in AD disease pathology.

The docking simulation process was carried out using PyDock, a rigid-body docking algorithm that incorporates electrostatics and desolvation energy scoring to predict protein-protein interactions (Jiménez-García et al., 2013). PyDock generates a large number of potential docking conformations and ranks them based on their binding energy scores, allowing for the identification of the most favorable interactions. In this study, a total of 10,000 docking simulations were performed, with each simulation producing an energy score that reflects the stability and favorability of the predicted ligand-receptor interaction. The energy score integrates multiple factors, including electrostatic complementarity, van der Waals interactions, and desolvation effects, providing a comprehensive evaluation of the docking pose. Lower energy scores correspond to more stable and potentially biologically relevant interactions. To visualize the distribution of docking energy scores across all simulations, a histogram was generated using Matplotlib’s hist function. The histogram was created with a bin size of 70 (bins=70), ensuring a detailed representation of the energy score distribution. This visualization provided insights into the overall quality of docking predictions, highlighting whether the docking results followed a normal distribution or exhibited clusters of particularly stable binding conformations.

## Results

### scRNA data background and annotation

While there are many integrated brain-linked cell atlas datasets in the published literature, appropriate datasets capturing sufficient numbers of neurons and also NVU component cells that can be used for CCC inference are limited. After vetting 31 papers (Sup Table S1), we selected and analysed murine brain scRNASeq data published from four independent papers relating to ageing, T2D and AD to identify common or contrasting gene expression patterns relating to CCC. The ageing groups were mice at 3, 18, and 24 months old. The chronological age of the mice in the AD groups was 6 or 7 months old. The age of the db/db mouse (T2D) group was 4 months old.

In these datasets, we reanalysed them from the raw matrix level and reannotated as well. The commonly found cell types such were identified through typical gene signatures (Fig 2), such as SLC1A2 for astrocytes, CLDN5 for endothelial cells, CTSS for microglial cells, CALD1 for mural cells, SNAP25 and GRIA2 for neurons, and PLP1 for oligodendrocytes. Other less abundant cell types in the brain were classified as neuroblasts (SOX4), epithelial cells (TTR), macrophages (HLA-E) while eppendymal cells were identified through expression of cilia-related genes such as RSPH1 and CCDC39. For easy reference, the study refers to the AD 6 and 7 month sets as AD6 and AD7, respectively (Fig 2). For the ageing group, there was in total 6707 cells while the number of cells decreased sharply according to chronological age, most notable in the 24-month group (3m:3218, 18m:2811, 24m:678) (Fig 2 A). The AD7 had more cells than other datasets selected with control group (16258 cells) and AD (25708 cells) (Fig 2 D). This was a key reason why it was added it into the study despite it having a 3 months time difference compared with the T2D dataset. On the other hand, AD6 only had 4992 cells in the AD condition which was less than half of its control (11049) (Fig 2 C). The non-T2D (control) mouse brain had less cells (3307) than the brain from T2D (4769 cells) (Fig 2 B).

**Figure 2.**
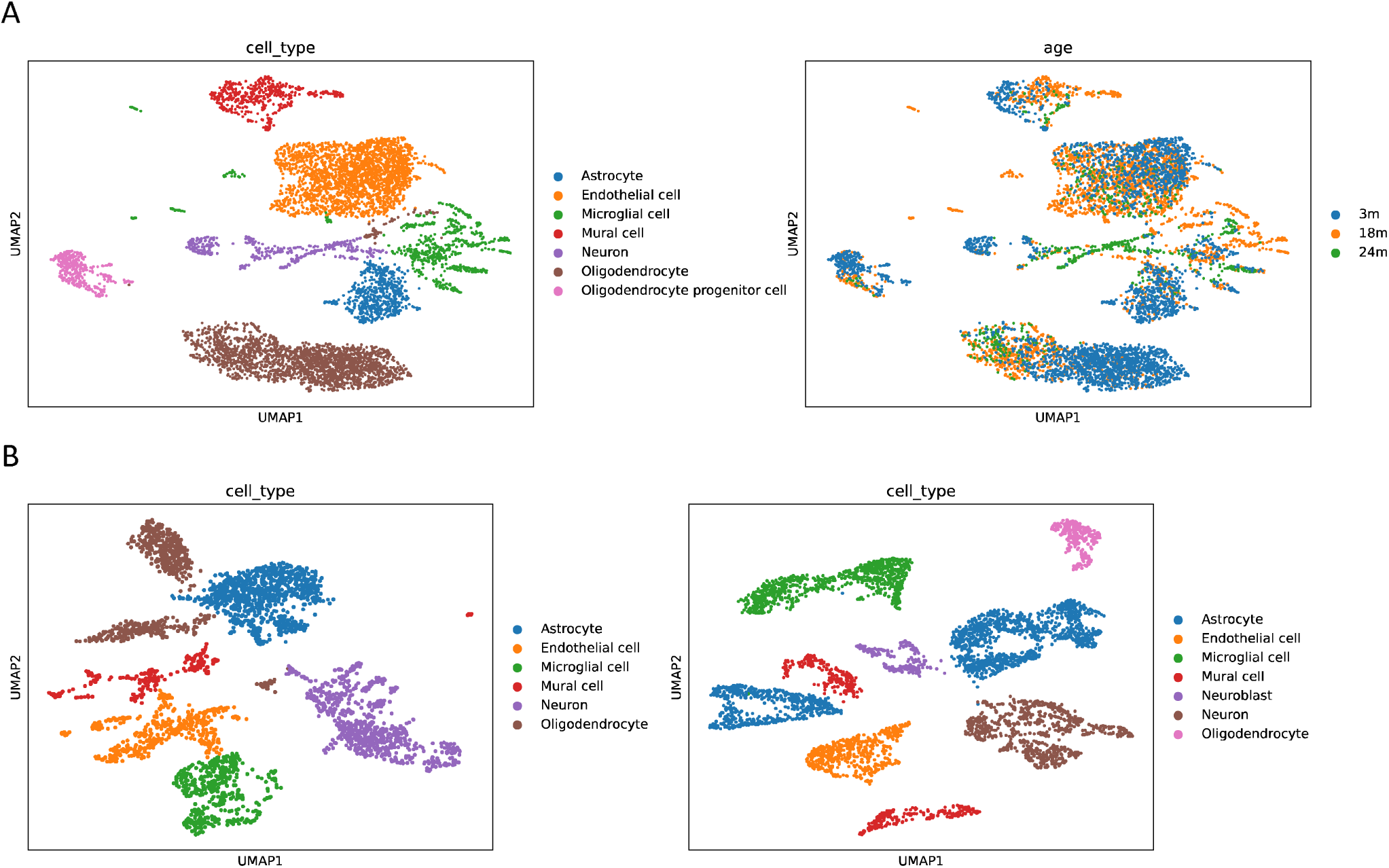
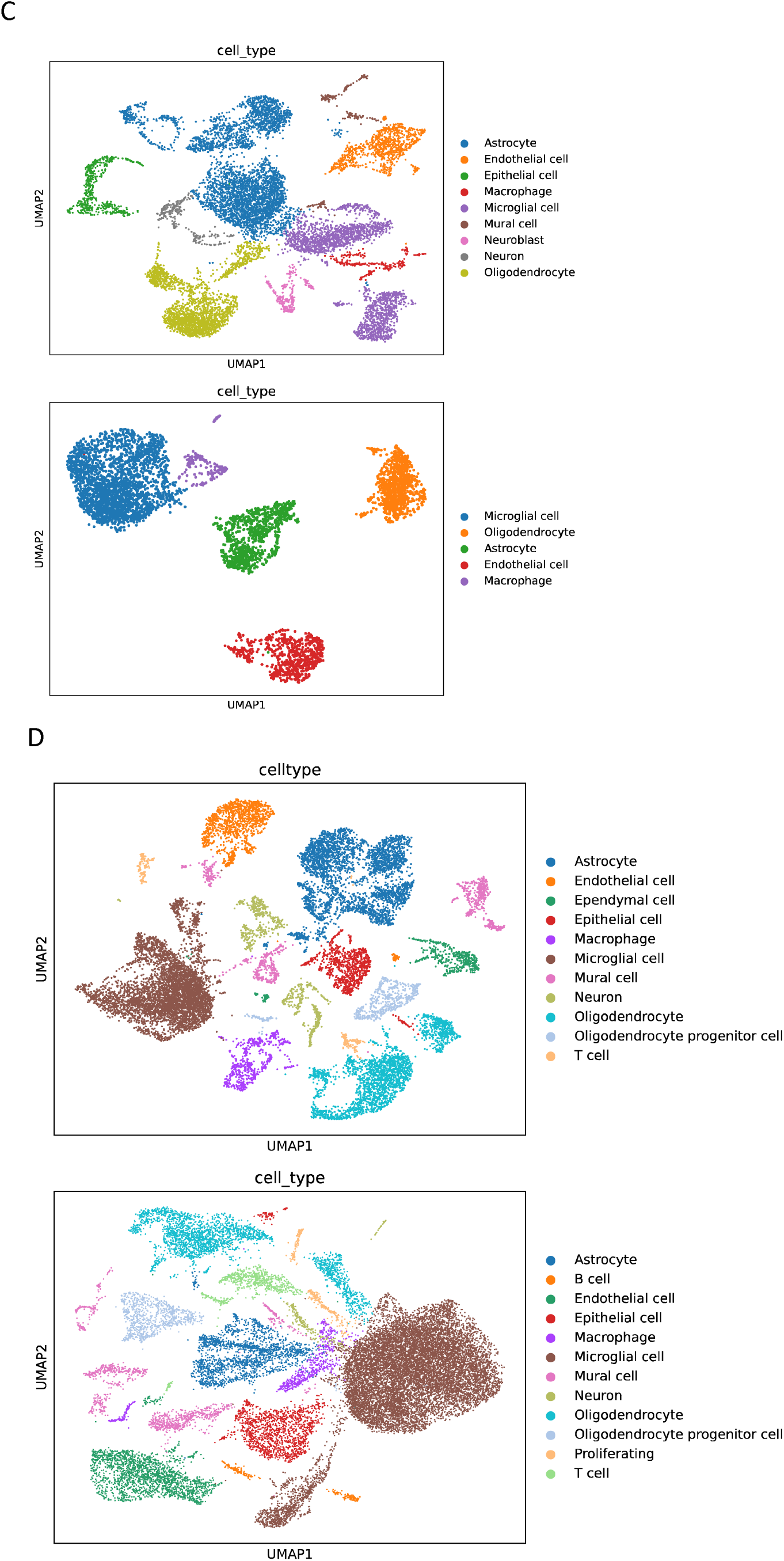
scRNA UMAP projections on four datasets. (A) Ageing dataset: left panel shows clustering by annotated cell types; right panel shows distribution of cells coloured by animal age group. (B) Type 2 Diabetes (T2D) dataset: left panel represents control samples; right panel represents cells from T2D samples. (C) Alzheimer’s disease dataset AD_6: upper panel displays control samples; lower panel shows AD cases. (D) Alzheimer’s disease dataset AD_7: upper panel shows control group; lower panel shows AD group. Across all datasets, each dot corresponds to a single cell, positioned according to transcriptional similarity in UMAP space, with colour schemes specific to each panel to highlight the relevant grouping variable.

### Ageing and AD featured loss of CCC in the brain while T2D showed gained propensity

The LIANA package (Dimitrov et al., 2022) was able to infer CCC in each brain dataset, after which an established comparison analysis (REF) was conducted to evaluate the impact on CCC. Among the four groups, ageing, AD6, and AD7 showed predominance of consensus and loss CCC although there were gained CCC in the T2D group. However, based on the categorical change count, the AD7 was more similar with ageing process than AD6 (Fig 3 A). On the other hand, the total number of changes in the AD6 and T2D group was less than the other two groups (ageing:1640, AD6:173, AD7:1874, T2D:368) (Fig 3 A). Even after normalizing by the total number of types of communication, AD6 and T2D changes were still lower than the other two (ageing: 33.469, AD6: 6.92, AD7: 15.616, T2D: 10.514).

**Figure 3.**
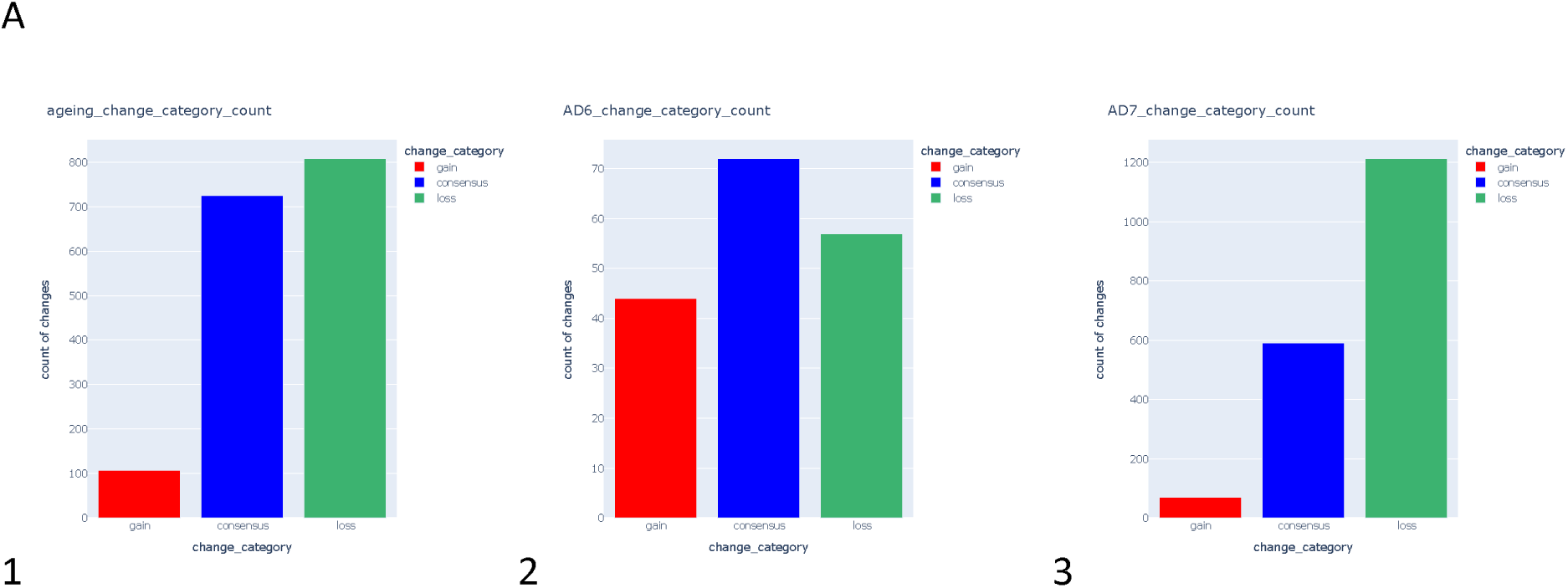
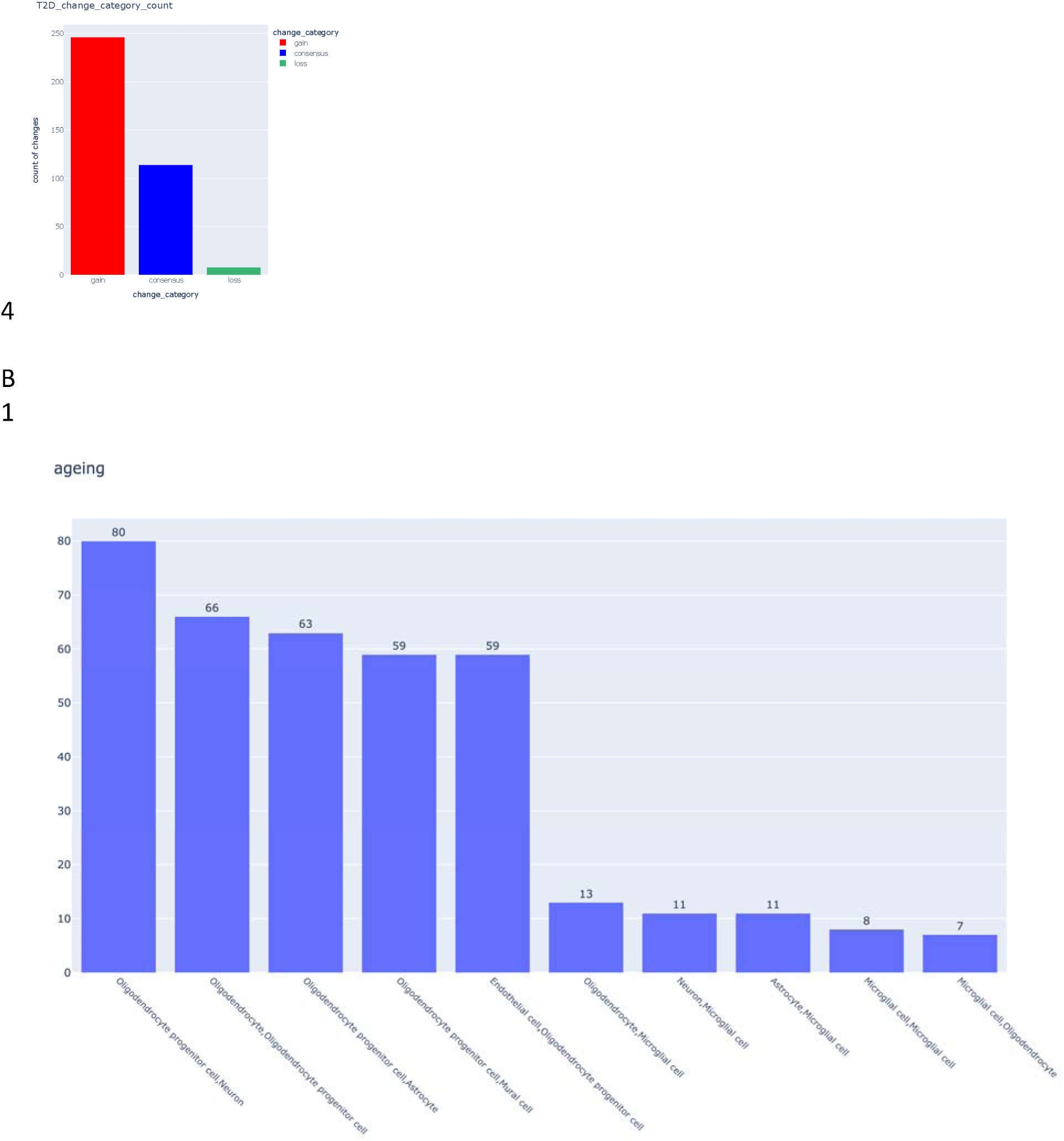
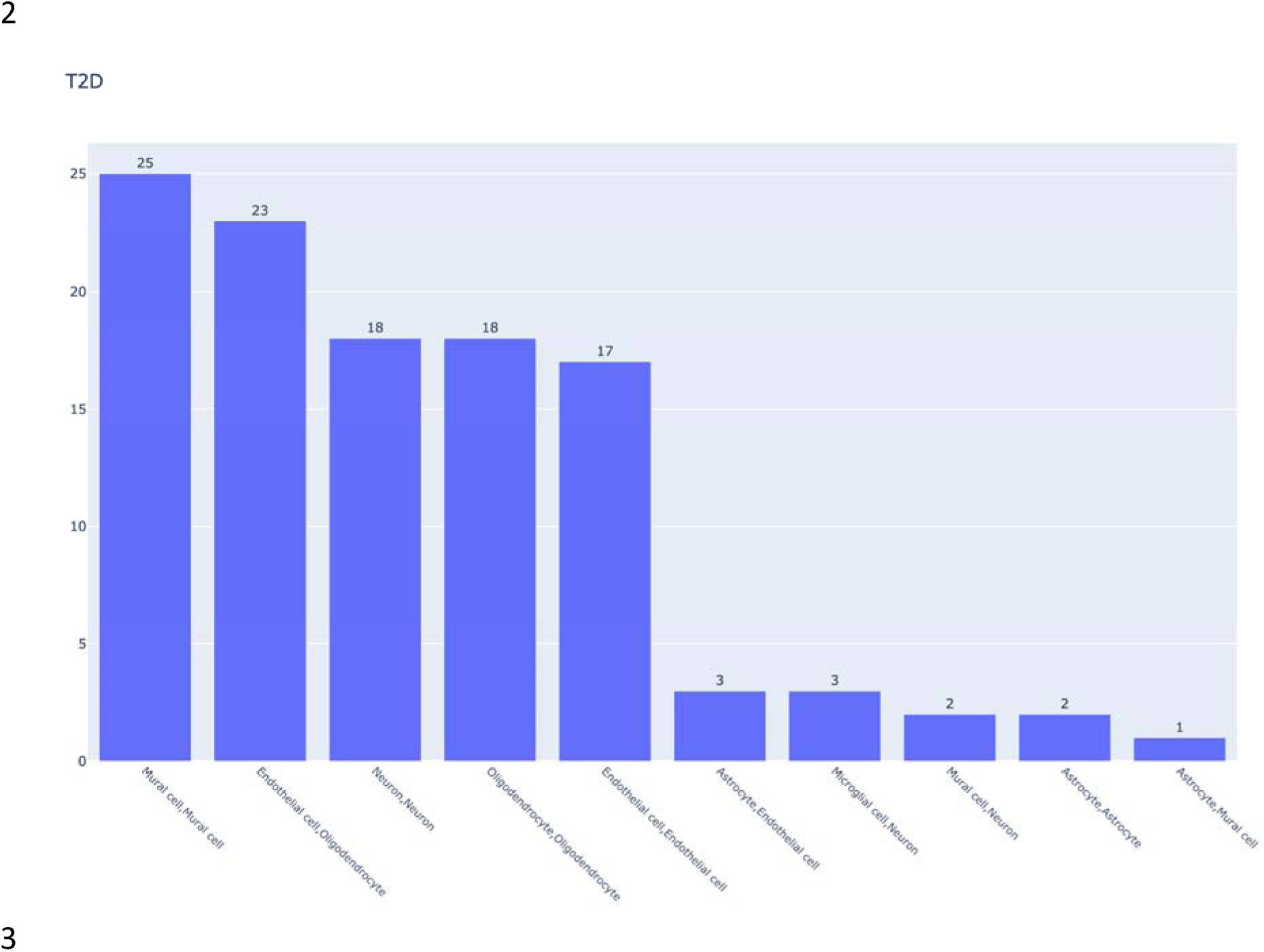
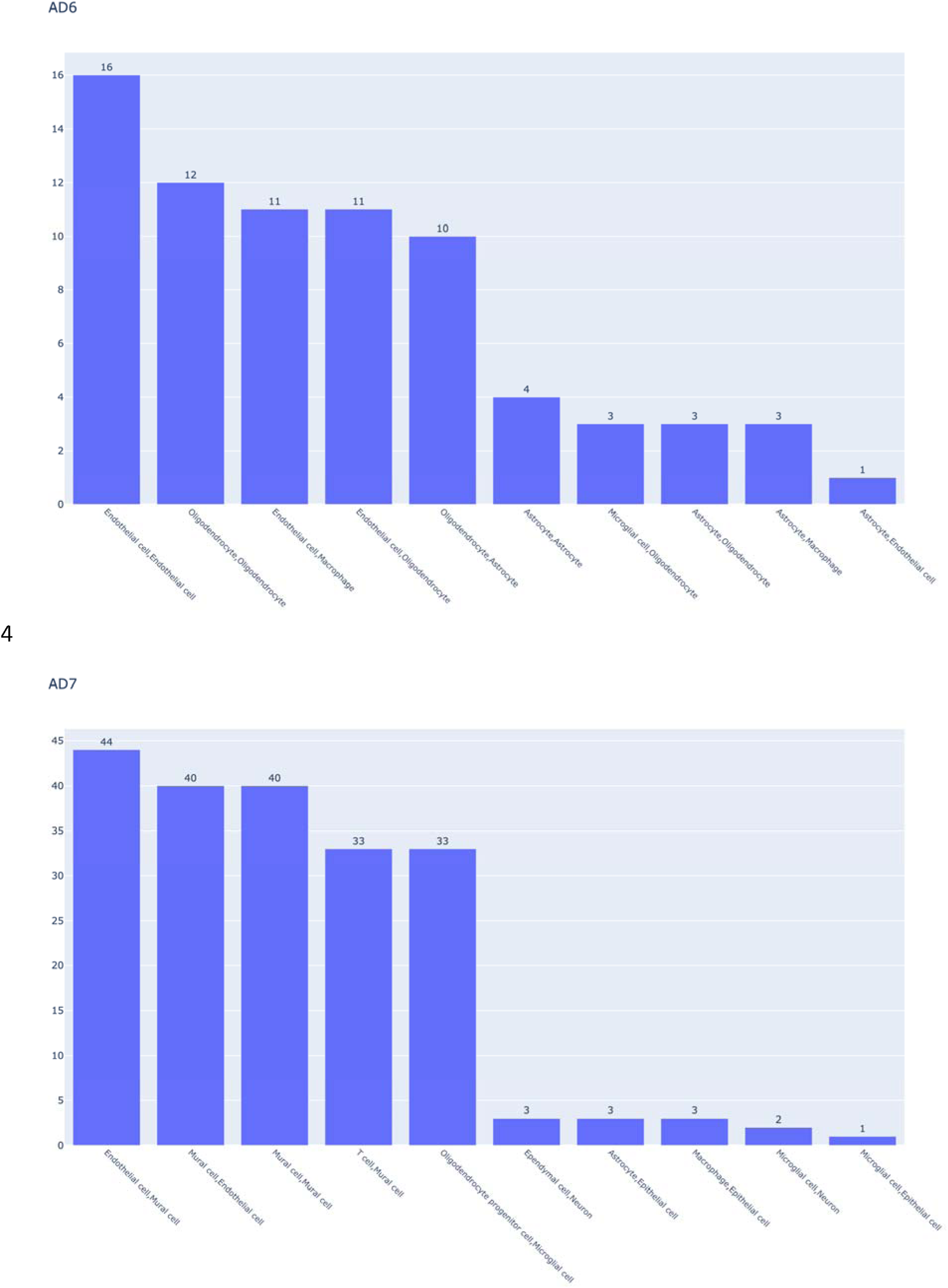
Cell to cell communication change between conditions in each dataset. Gain refers to newly established ligand receptor pairs while loss meant lost ligand receptor pairs. Consensus indicated the number of ligand receptpr pairs that existed in both conditions but the communication strength was significantly changed. (A) bar chart counting the number of categorical changes by datasets. This highlighted the featured CCC change category in each dataset. (B) the types of communication in each dataset ranked by the total number of changes regardless of change category (only the top 5 and the bottom 5 were shown here, to see the full list, check Sup Fig 1). The bar indicated the magnitude of change for each type of communication.

With ageing, ‘Oligodendrocyte progenitor cell,Neuron’ had the highest number of CCC changes (80), followed by ‘Oligodendrocyte,Oligodendrocyte progenitor cell’ (66), ‘Oligodendrocyte progenitor cell,Astrocyte’ (63), ‘Oligodendrocyte progenitor cell,Mural cell’ (59), ‘Endothelial cell,Oligodendrocyte progenitor cell’ (59) (Fig 3 B1). Thus, oligodendrocytes and oligodendrocyte progenitor cells appear to be the most important cell types showing altered CCC with chronological age.

With respect to AD, there was an interesting discovery showing the importance of endothelial CCC. Homotypic ‘Endothelial cell,Endothelial cell’ CCC was the top ranked with 16 changes in the AD6 group (Fig 3 B3). The third and fourth were also endothelial cell related types of CCC. Furthermore, the AD7 group also manifested vascular-associated CCC changes as four out of top five involved endothelial cells or mural cells (Fig 3 B4).

T2D brain scRNASeq datasets showed marked differences when compared with ageing datasets. When non-T2D and T2D brain CCC changes were assessed, the top 5 highest levels were ‘Mural cell,Mural cell’ (25), ‘Endothelial cell,Oligodendrocyte’ (23), ‘Neuron,Neuron’ (18), ‘Oligodendrocyte,Oligodendrocyte’ (18), ‘Endothelial cell,Endothelial cell’ (17) (Fig 3 B2), which showed a stronger vascular cell types association compared with ageing, but not as much as AD.

### Age-related diseases featured vascular changes compared with ageing changes

Next, we investigated how cell types changed in terms of expressing or receiving CCC signals. It was found that Oligodendrocyte progenitor cells had the strongest change for ligands as the source cell type while neurons showed the highest receptor change pattern as a target cell type in the ageing group (Sup Fig 2). Microglial cells maintained the lowest rank in both source and target cell types (Sup Fig 2). In the T2D group, endothelial cells were ranked as the top source cell type and mural cells as the top target cell type, thus highlighting the importance of these cells to the integrity of the NVU (Sup Fig 2).

In the AD6 group, endothelial cells were the cell type most changed for both source and target (Sup Fig 2). On the other hand, in the AD7 group, mural cells were the highest ranked cell type, for both source and target (Sup Fig 2). However, there was no mural cell annotated in AD6. Therefore, comparison on mural cell between the AD6 and AD7 groups was not possible.

When this data is taken together, there is a strong suggestion that changes in brain endothelial and mural cell CCC are of critical importance in AD and T2D when compared to normal healthy ageing.

### Several key ligand receptor pairs were identified as key common CCC targets in AD and T2D

To find common CCC factors behind ageing and T2D and AD, the next step taken was to screen out some key changes contributing to ageing and these two age-linked diseases. The comparison between AD6 and T2D is referred to as AD6-T2D and comparison between AD7 and T2D as AD7-T2D. The AD6-T2D had less overlapping changes (100) than AD7-T2D (258) which was anticipated due to the lower number of cell types (Fig 4 A). However, the ratio of overlapping changes was higher (100/258, ∼ 0.38759) than the ratio by types of communication (AD6/AD7, 25/120, ∼ 0.2083) which is suggestive of convergence on common factors between AD and T2D.

**Figure 4.**
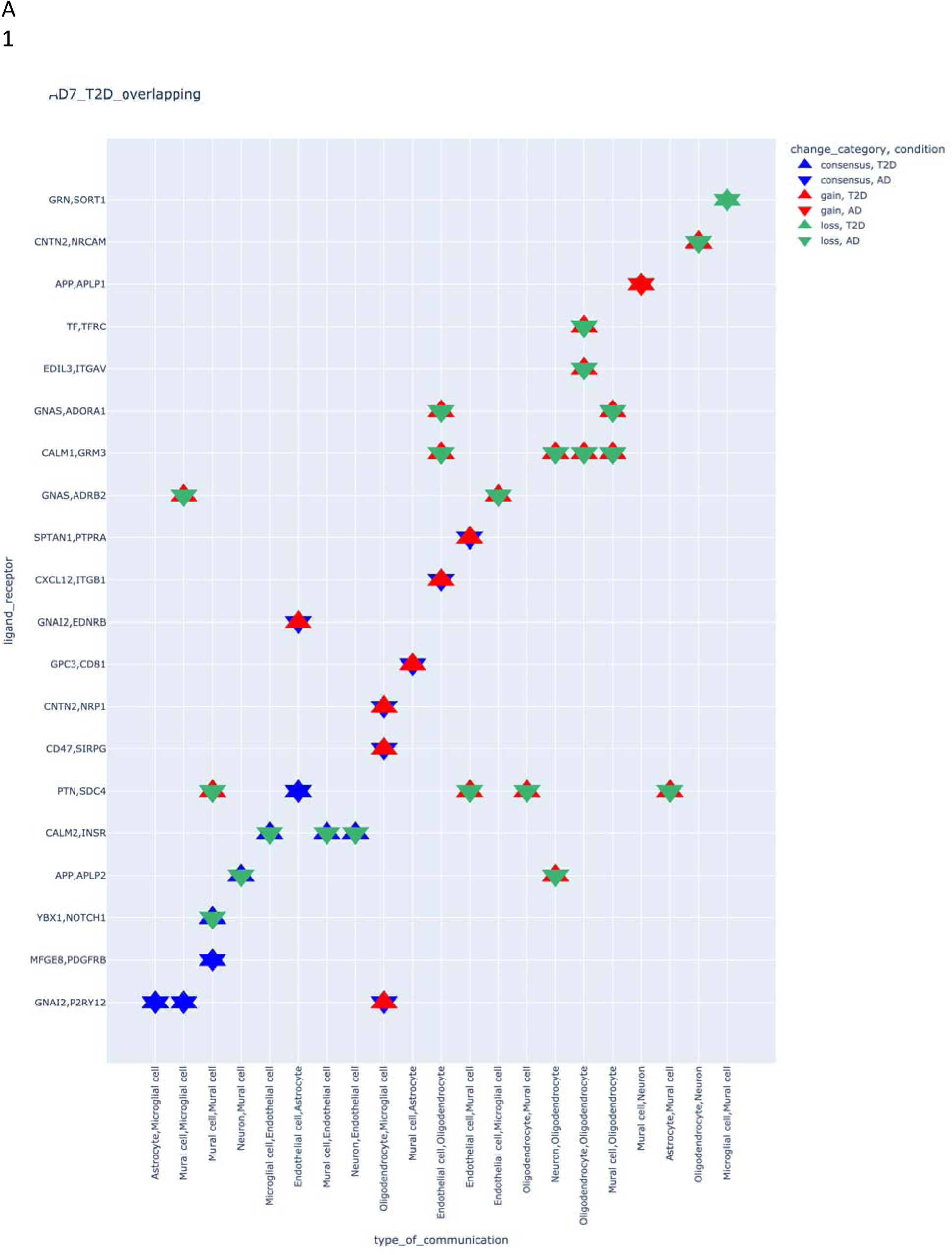
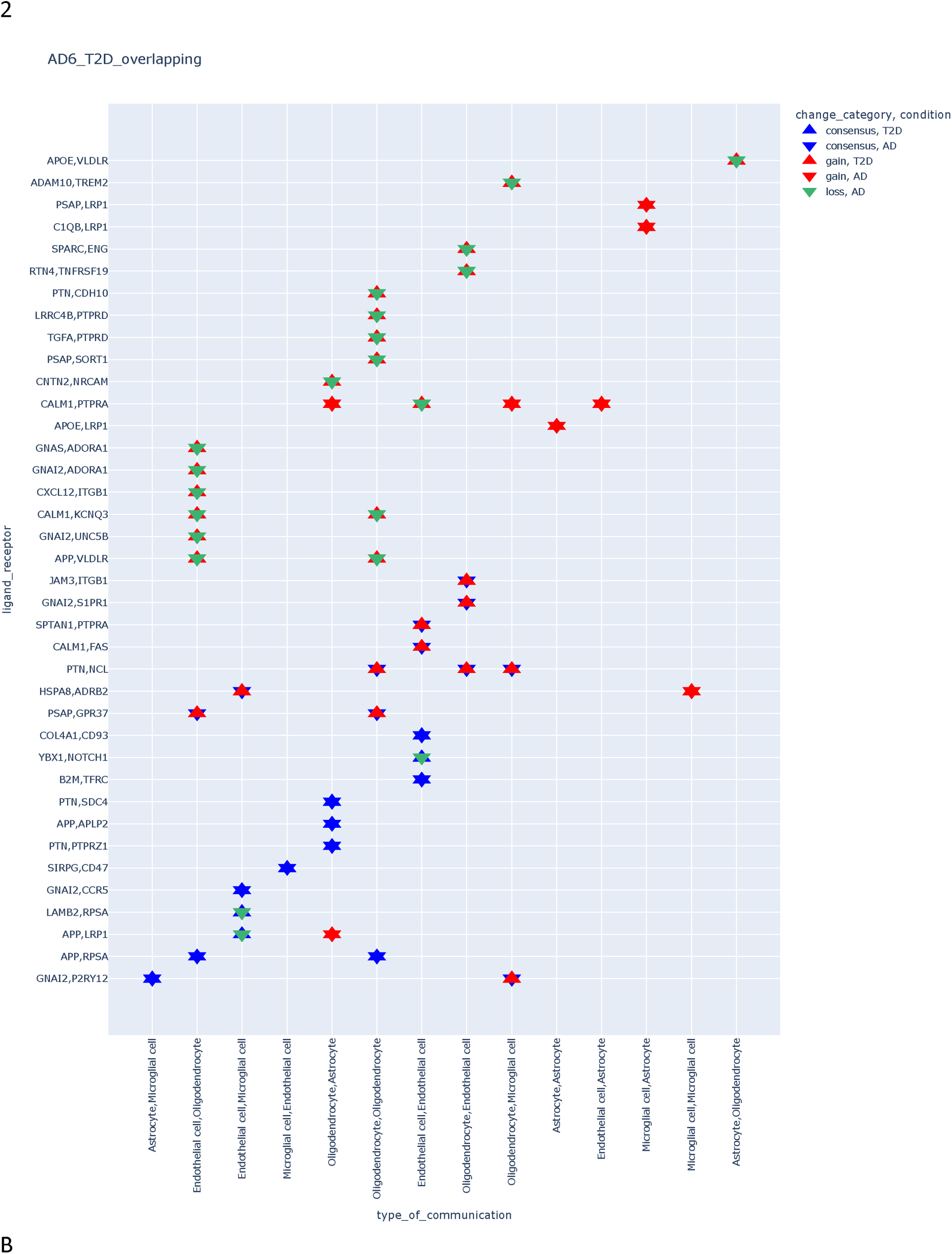
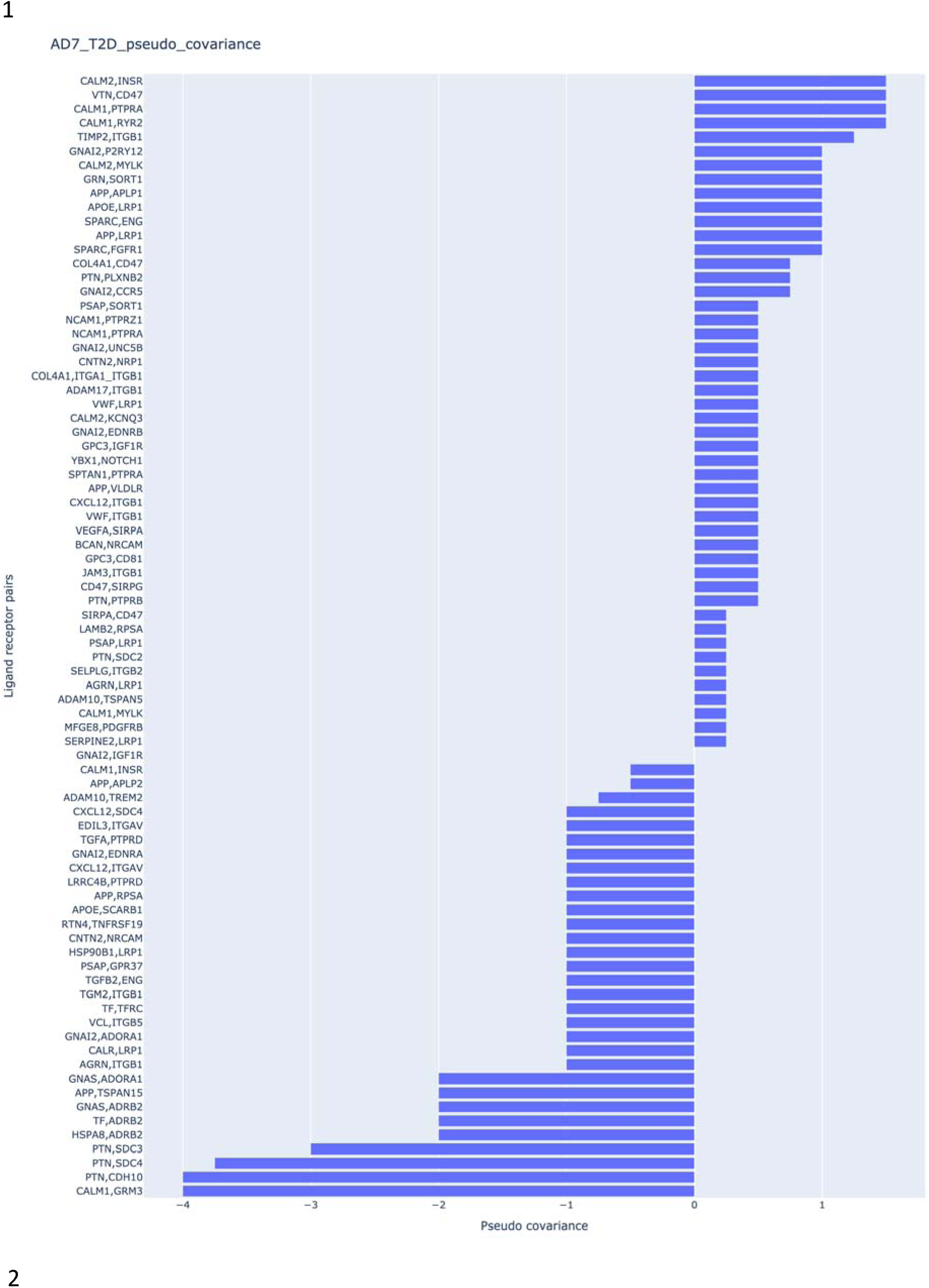
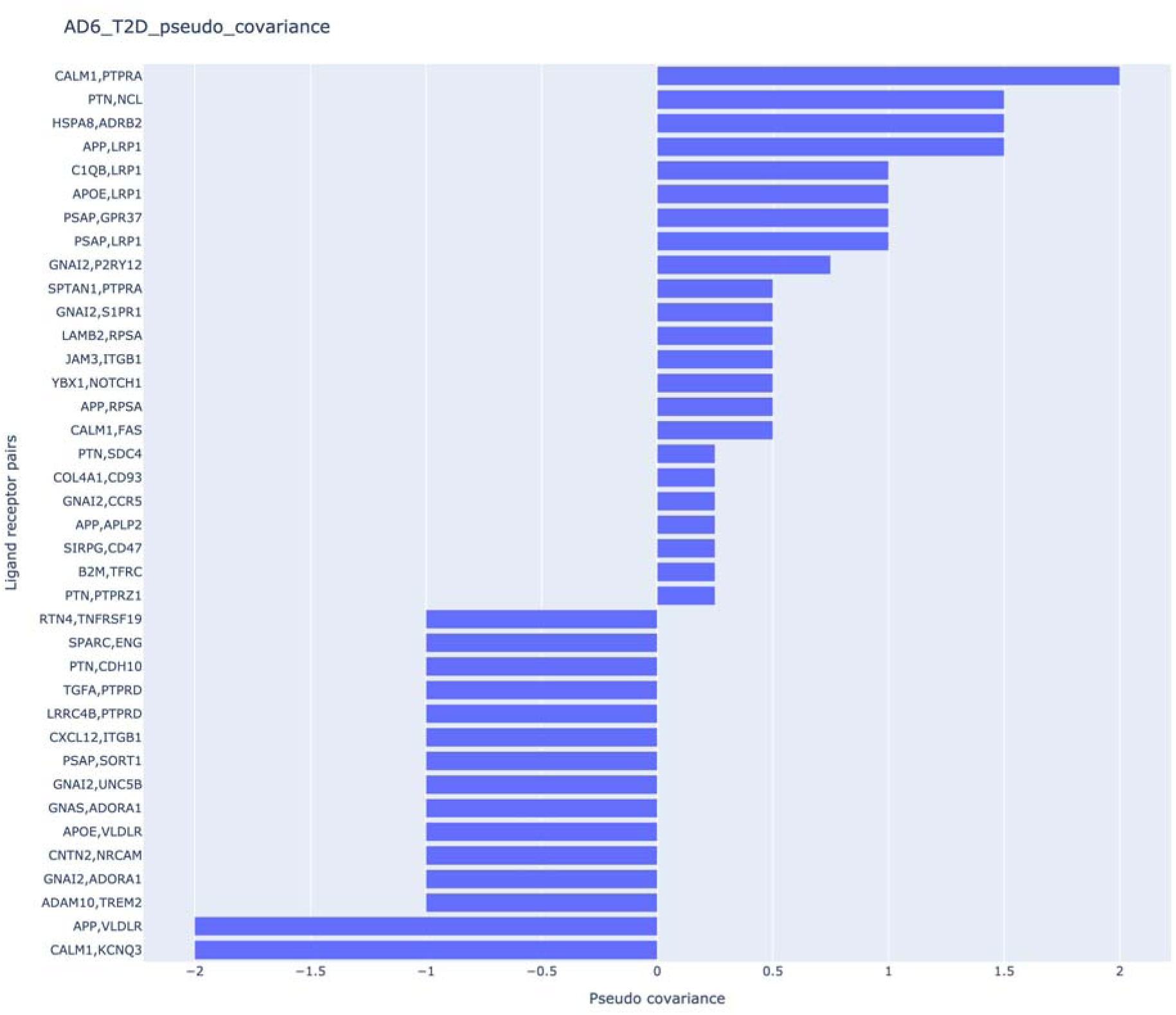
(A) Overlapping parts changes between AD7 and T2D (1) only part of the whole set is shown here for visualization, the full set is available in Sup Table S3, or AD6 and T2D (2, full set). The same colour in the same coordinate indicated convergence. Up triangle refers to T2D condition while down triangle is for AD. (B) pseudo-covariance bar chart of ligand receptor paired ranked between AD_7 and T2D (1), or AD_6 and T2D (2). The high convergence ligand receptor pairs would be ranked as high positive values.

The most overlapped ligand receptor pairs in AD7-T2D were HSP90B1, LRP1, PTN, SDC4, PTN, and CDH10 while the counterparts in AD6-T2D included CALM1, PTPRA, PTN, NCL, APP, and VLDLR (Fig 4 A, Sup Table S3). Some ligands had a frequent overlap between AD6 and AD7, such as PTN, APP, and GNAI2. However, many of these overlapping ligands did not show the same direction changes between T2D and AD, indeed, they often revealed opposing CCC profiles in terms of loss vs gain. In the AD6-T2D comparison, 18 out of 50 overlapping parts were in the opposite direction while a slightly higher proportion was observed in the AD7-T2D comparison (51 out of 129).

To identify ligand-receptor pairs, a pseudo covariance formula (see methods) was established to sort the divergence and convergence between T2D and AD conditions. The formula assumed the opposite direction as a negative value to the pseudo covariance, a positive value if consensus direction was involved, and at last the highest positive value if the directions were double loss or double gain. Hence, the ligand-receptor pairs were sorted and ranked by their pseudo covariance values from positive to negative.

Within the top 10 positive covariance ligand receptor pairs from AD_7_T2D and AD_6_T2D (Fig 4 B), three pairs were shared: ‘APP,LRP1’, ‘CALM1,PTPRA’, ‘GNAI2,P2RY12’ (Fig 4 B). These three pairs appear to show commonality of CCC in the brain of mice with T2D and AD and warrant further investigation.

### LRP1 receptor as a potential driver in brain pathology associated with ageing, T2D and AD

From the global gene expression perspective, ageing-related changes predominantly led to a loss of CCC (Fig 5 A). However, between the age ranges of 3–18 months and 18–24 months, both gain and loss of CCC were observed, creating conflicting patterns. These conflicts were deemed invalid by the algorithm, and we assumed that CCC changes only found in pathological conditions should be corrected by default (see methods). Notably, certain molecules—PTN, NCL, APOE, LRP1, GNAI2, P2RY12, APP, and APLP1—exhibited particularly complex ageing-related patterns (Fig 5 A). After computing the correction actions, the algorithm identified that the primary adjustments involved microglial cells, oligodendrocytes, endothelial cells, mural cells, and astrocytes (Table 1.1, Table 1.2). One key finding was that the action “decrease LRP1 from astrocytes” was promoted three times, derived from both AD6-T2D-ageing and AD7-T2D-ageing datasets. This suggests that LRP1 in astrocytes plays a crucial role as an age-related factor contributing to disease progression in the brain.

**Figure 5.**
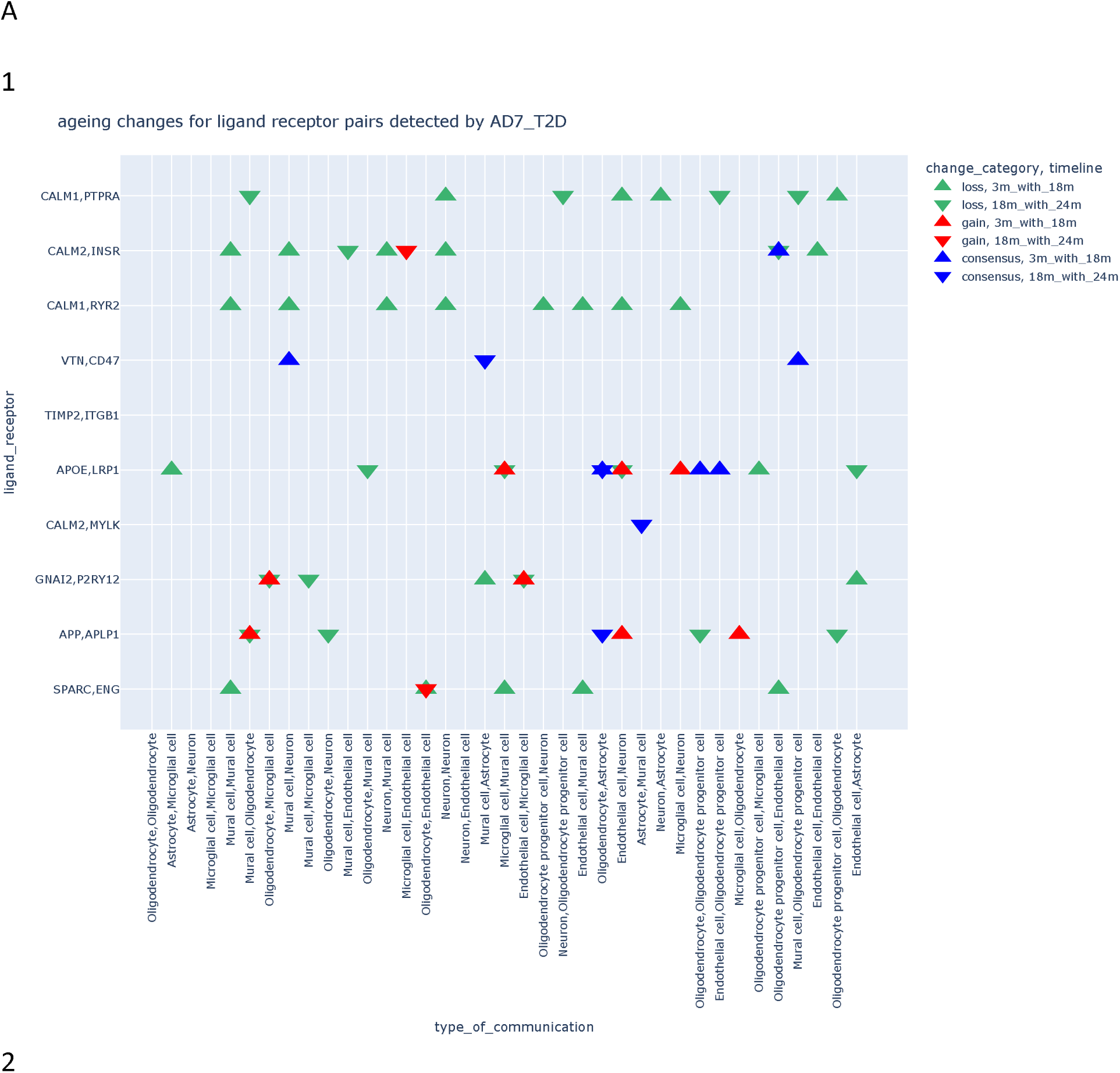
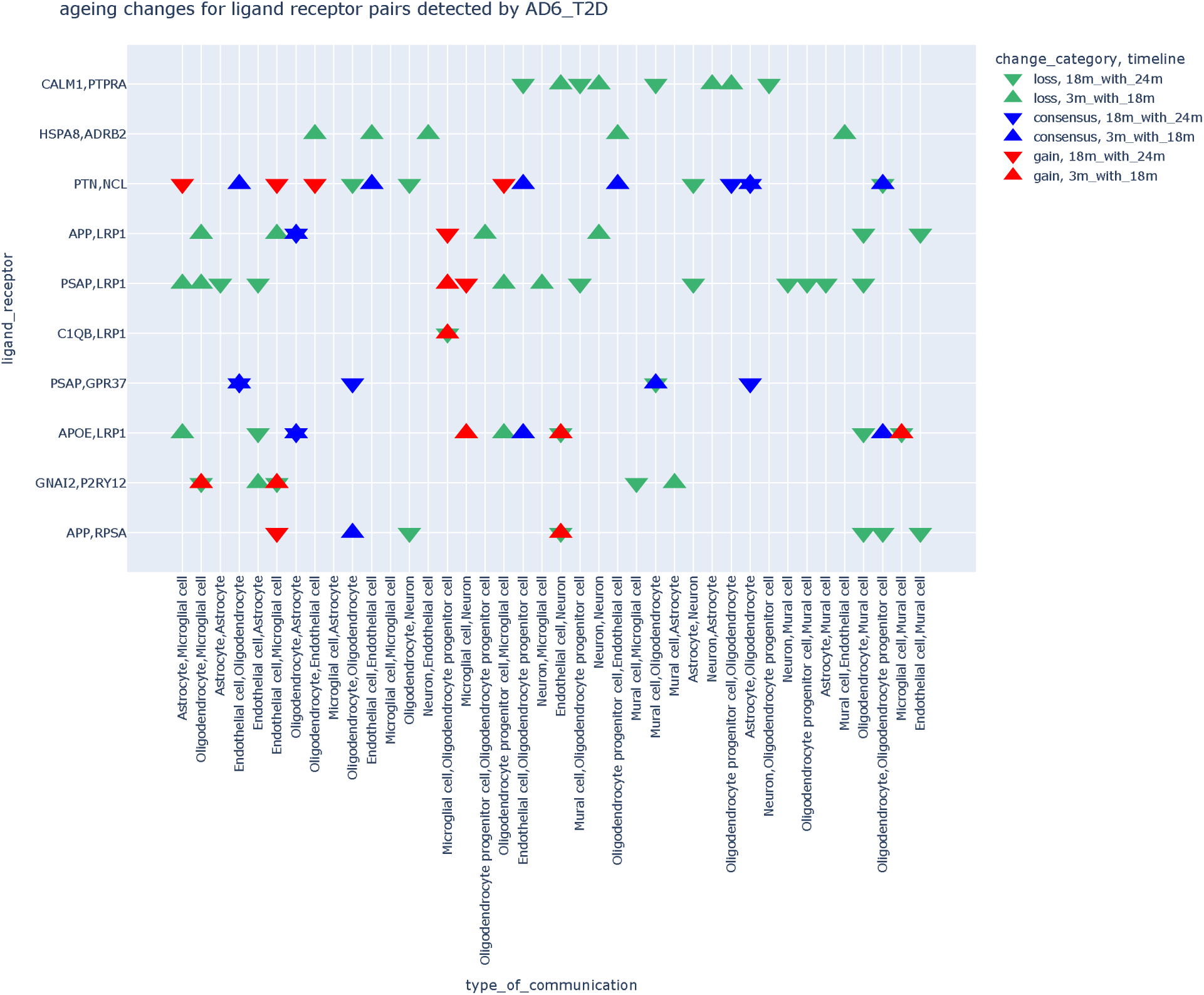
(A) The ageing change situation of the ligand receptor pairs that had high pseudo covariance in (1) AD_7_T2D (2) AD_6_T2D. Table1 prevention action result for AD_6_T2D_ageing_action (see methods), Table2 prevention action result for AD_7_T2D_ageing_action.

**Table 1.1.**

| AD_6_T2D_ageing_action | Gene Action | Affected Cell Types |
| --- | --- | --- |
| 0 | decrease_CALM1_from | Oligodendrocyte |
| 1 | decrease_PTPRA_from | Microglial cell |
| 2 | improve_CALM1_from | (empty) |
| 3 | improve_PTPRA_from | (empty) |
| 4 | decrease_HSPA8_from | Microglial cell |
| 5 | decrease_ADRB2_from | Microglial cell |
| 6 | improve_HSPA8_from | (empty) |
| 7 | improve_ADRB2_from | (empty) |
| 8 | decrease_APP_from | (empty) |
| 9 | decrease_LRP1_from | Astrocyte |
| 10 | improve_APP_from | Endothelial cell |
| 11 | improve_LRP1_from | Microglial cell |
| 12 | decrease_PTN_from | (empty) |
| 13 | decrease_NCL_from | Endothelial cell, Microglial cell |
| 14 | improve_PTN_from | (empty) |
| 15 | improve_NCL_from | (empty) |
| 16 | decrease_APOE_from | (empty) |
| 17 | decrease_LRP1_from | (empty) |
| 18 | improve_APOE_from | (empty) |
| 19 | improve_LRP1_from | (empty) |
| 20 | decrease_C1QB_from | Microglial cell |
| 21 | decrease_LRP1_from | Astrocyte |
| 22 | improve_C1QB_from | (empty) |
| 23 | improve_LRP1_from | (empty) |
| 24 | decrease_PSAP_from | Oligodendrocyte, Endothelial cell |
| 25 | decrease_GPR37_from | (empty) |
| 26 | improve_PSAP_from | (empty) |
| 27 | improve_GPR37_from | (empty) |
| 28 | decrease_PSAP_from | Microglial cell |
| 29 | decrease_LRP1_from | (empty) |
| 30 | improve_PSAP_from | (empty) |
| 31 | improve_LRP1_from | (empty) |
| 32 | decrease_GNAI2_from | Oligodendrocyte |
| 33 | decrease_P2RY12_from | (empty) |
| 34 | improve_GNAI2_from | (empty) |
| 35 | improve_P2RY12_from | (empty) |
| 36 | decrease_SPTAN1_from | (empty) |
| 37 | decrease_PTPRA_from | Endothelial cell |
| 38 | improve_SPTAN1_from | (empty) |
| 39 | improve_PTPRA_from | (empty) |

**Table 1.2.**

| AD_7_T2D_ageing_action | Gene Action | Affected Cell Types |
| --- | --- | --- |
| 0 | decrease_CALM1_from | Oligodendrocyte |
| 1 | decrease_PTPRA_from | Mural cell, Microglial cell |
| 2 | improve_CALM1_from | (empty) |
| 3 | improve_PTPRA_from | (empty) |
| 4 | decrease_CALM2_from | (empty) |
| 5 | decrease_INSR_from | (empty) |
| 6 | improve_CALM2_from | Neuron, Mural cell |
| 7 | improve_INSR_from | (empty) |
| 8 | decrease_CALM1_from | Oligodendrocyte, Astrocyte |
| 9 | decrease_RYR2_from | (empty) |
| 10 | improve_CALM1_from | (empty) |
| 11 | improve_RYR2_from | (empty) |
| 12 | decrease_VTN_from | Mural cell |
| 13 | decrease_CD47_from | Astrocyte, Oligodendrocyte, Mural cell |
| 14 | improve_VTN_from | (empty) |
| 15 | improve_CD47_from | (empty) |
| 16 | decrease_TIMP2_from | Oligodendrocyte, Mural cell |
| 17 | decrease_ITGB1_from | Mural cell |
| 18 | improve_TIMP2_from | (empty) |
| 19 | improve_ITGB1_from | (empty) |
| 20 | decrease_CALM2_from | Oligodendrocyte |
| 21 | decrease_MYLK_from | Mural cell |
| 22 | improve_CALM2_from | (empty) |
| 23 | improve_MYLK_from | (empty) |
| 24 | decrease_APP_from | (empty) |
| 25 | decrease_LRP1_from | Astrocyte |
| 26 | improve_APP_from | Neuron |
| 27 | improve_LRP1_from | Mural cell |
| 28 | decrease_GNAI2_from | Oligodendrocyte |
| 29 | decrease_P2RY12_from | (empty) |
| 30 | improve_GNAI2_from | (empty) |
| 31 | improve_P2RY12_from | (empty) |
| 32 | decrease_APP_from | Mural cell |
| 33 | decrease_APLP1_from | (empty) |
| 34 | improve_APP_from | (empty) |
| 35 | improve_APLP1_from | (empty) |
| 36 | decrease_SPARC_from | (empty) |
| 37 | decrease_FGFR1_from | (empty) |
| 38 | improve_SPARC_from | (empty) |
| 39 | improve_FGFR1_from | (empty) |

Finally, the current study also attempted to incorporate resveratrol treatment as a positive reference, based on the evidence that this molecule has neuroprotective properties, including improved blood blow, anti-inflammatory effects, and clearance of amyloid-beta (AB) (Bastianetto et al., 2015; Gu et al., 2021; Santos et al., 2016; Yuan et al., 2024; Zhou et al., 2021). This bulk RNA sequence data was derived from a published study based on healthy mice injected with resveratrol (Khoury et al., 2018). This analysis showed several key genes that coincide with the CCC data, most notably within the top 10 positively correlated overlapping parts including CD47, LRP1, and FGFR1 from AD_7, only LRP1 from AD_6.

This CCC analysis combined with other strands of transcriptomics data, reveals that LRP1 could play an important modulatory role in brain pathology and could be a potential target for prevention of age-related brain changes, an *in silico*-based finding that warrants further mechanistic investigation.

### Binding affinity investigation of LRP1 ligands

To investigate how LRP1 interacts with key molecules involved in CCC, the study extracted protein structure (PDB) files from the RCSB Protein Data Bank for several ligands and receptors (see methods). These molecular structures were then subjected to docking simulations using the PyDock web tool (Fig 6 A), which generated 10,000 potential conformations for each ligand-receptor pair, along with binding energy scores for each conformation. This approach was then used to analyse five ligand-LRP1 interaction pairs relevant to the brain: PSAP-LRP1, APOE-LRP1, C1QB-LRP1, AB (Amyloid Beta)-LRP1 and APP-LRP1

**Figure 6.**
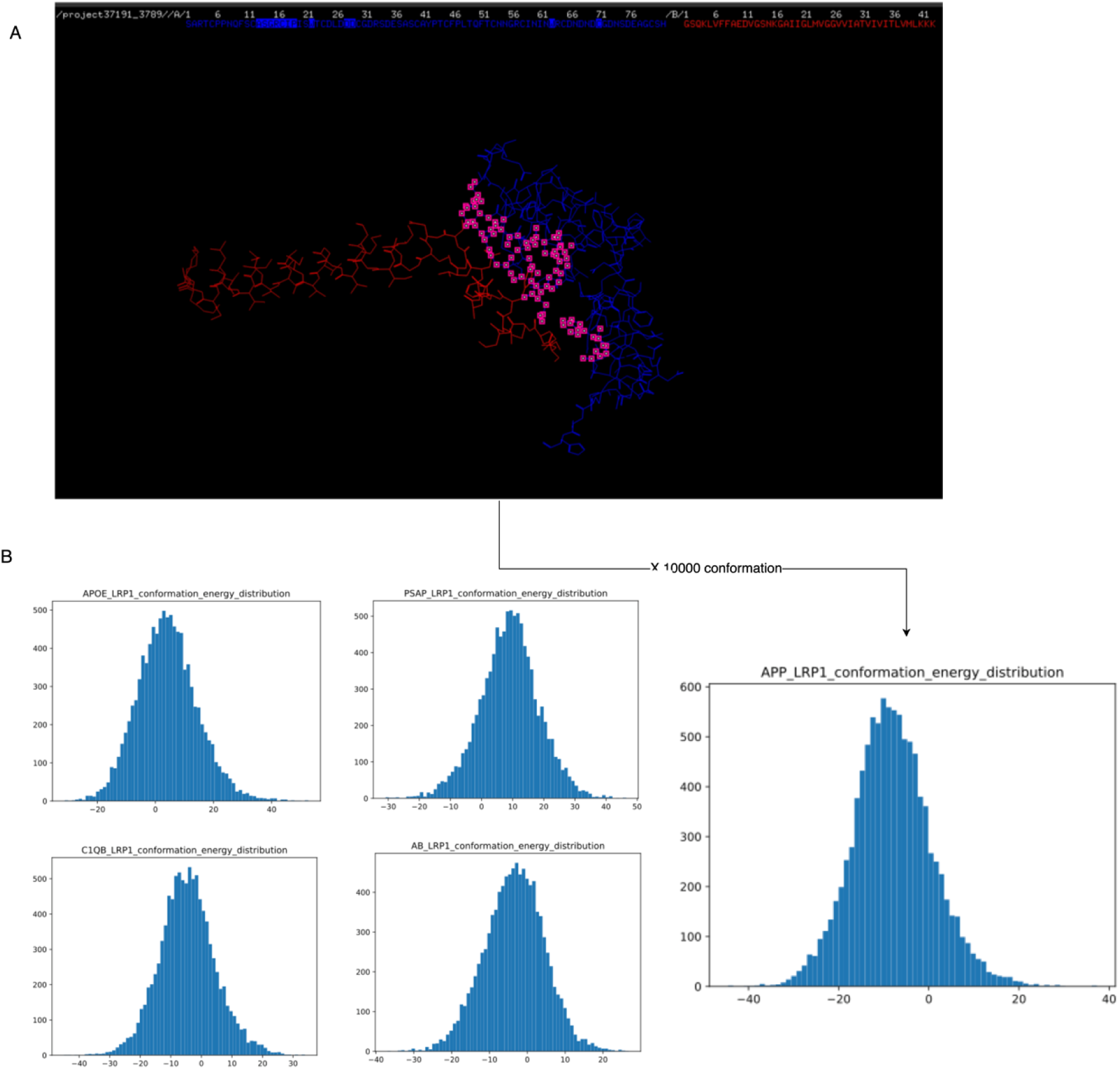
(A) Docking simulation conformation example generated from pymol. The blue structure is LRP1, the red structure is APP. Labelled regions on LRP1 represented the binding sites inference. (B) Energy score distribution by 10000 simulations.

### The importance of APP-LRP1 interaction as a high affinity CCC pairing

Energy score distributions revealed distinct binding propensities. PSAP and APOE interactions with LRP1 exhibited peaks above zero indicative of weak affinity and a low probability of stable binding (Fig 6 B), thus suggesting that these ligands are unlikely to contribute significantly to LRP1-mediated CCC in the brain. C1QB and AB showed more enhanced binding with LRP1 based on the energy distribution being were shifted leftward and suggestive of a stable interaction. APP-LRP1 demonstrated the strongest binding affinity, with its energy distribution peak positioned further left than C1QB and AB. This suggests that APP-LRP1 interaction is more important than other ligand receptor pair in terms of CCC associated with brain degenerative changes.

## Discussion

### Study design review

The current study has analysed four distinct scRNA-seq datasets, from the brain of ageing mice and compared this with matched datasets from murine models of AD and T2D. Our previously established model showed widely applicable capability to general two-conditions comparison by categorizing CCC changes into gain, consensus and loss.

A fundamental challenge in scRNA-seq studies involving multiple datasets is the trade-off between dataset integration and biological signal preservation. Integration techniques are often used to align datasets, enhancing the identification of shared cell types across conditions (REF). However, such integration can also obscure critical biological differences, potentially diminishing key pathological insights. In the context of CCC inference, this study prioritized a self-to-self comparison approach, where each condition was analysed independently without integration. This decision was based on the fact that CCC inference inherently does not favour integration, as it relies on within-condition ligand-receptor interactions rather than across-condition alignments. As a consequence of processing each dataset separately, there was scope to enable cell type composition and identity shifts to be identified and thus emerge as potential contributing factors in downstream analyses. This does not imply that the CCC results are unreliable but rather that the observed CCC changes could, in part, be influenced by shifts in cell populations (REF). This could be further explored through additional scRNA-seq analyses if needed although such secondary analyses was beyond the scope of this study. It is also important to acknowledge that transcriptomics-based approaches do not capture non-peptide mediators of intercellular communication (e.g., steroid hormones, metabolites) (Alberts et al., 2002). Therefore, small molecule communication is not represented in this framework. Also, as discussed in our previous CCC analysis across the life-span (REF), the ligand receptor pairings suggested here are not limited to typical biological ligand receptor pairs. Based on this non-biased, *in silico* analysis there are a number of unknown or unexpected interactions and these would need to be validated using further experimental approaches.

### Integrative analysis has identified LRP1 as a key factor linked to brain ageing and age-related disease

The CCC changes occurring in murine brain, as revealed in this study, suggest that changes related to AD are more age-linked than T2D. Indeed, both healthy ageing and AD featured overall loss of CCC events no matter whether comparing 6 or 7 months of AD duration (Fig 7). By contrast, the brain from T2D mice showed a gain in CCC events. There were, however, some commonalities and from all datasets analysed, there were clear vascular cell-associated CCC alterations in the brain which underscores the importance of these cells and the nature of cell communication for integrity of the NVU, especially in relation to AD and T2D.

**Figure 7.**
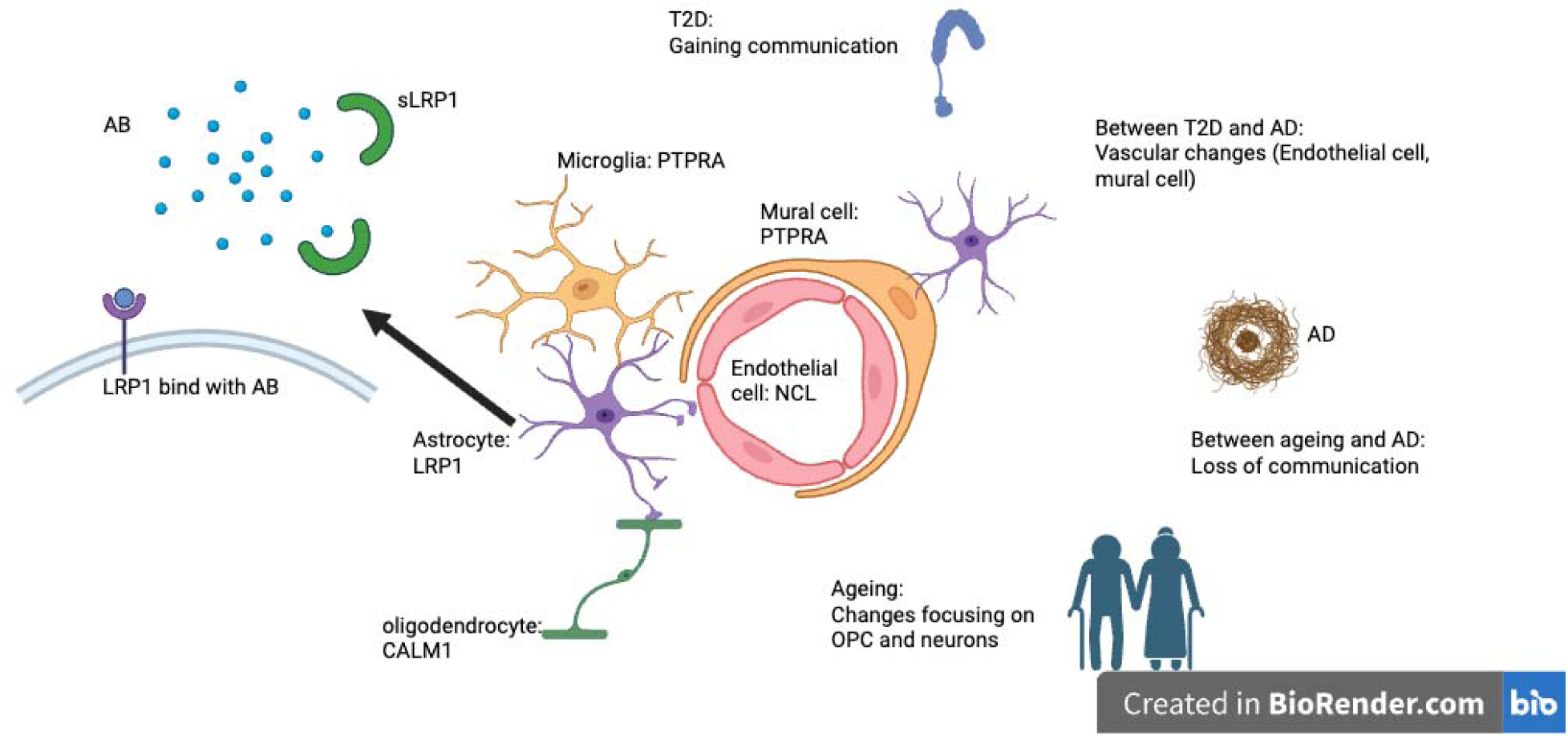
a diagram summary of findings from integrative analysis. T2D was represented by insulin receptor symbol while AD was represented by AB plaque. Central neurovascular unit showed some key molecules and cell types identified as common factors driving normal ageing into age-related diseases.

To uncover potential brain gene expression pathways which are common to ageing, AD and T2D, the overlapping CCC changes were further assessed. The overlapping parts dramatically narrowed down the range of ligand receptor pairs of interests. Despite a proportion of oppositely overlapped changes, some positively correlated ligand receptor pairs were revealed by pseudo covariance analysis. These included as ‘APP,LRP1’, ‘CALM1,PTPRA’, and ‘GNAI2,P2RY12’. Although both AD6-T2D and AD7-T2D exhibited some calmodulin-related CCC, it should be noted that calmodulin is regarded as an intracellular signalling molecule although some studies have suggested that it can be secreted by cells (O’Day et al., 2012). Similarly, PTPRA has been mainly described as an intracellular signalling transduction molecule although there is some evidence suggesting its role in extracellular communication (Yadav et al., 2020; Yao et al., 2017) and it has been found to be essential for nervous system development such as neuronal cell migration and synaptic plasticity (Petrone et al., 2003). The potential importance of PTPRA in brain degenerative pathology might depend on the communication with CALM1 but this requires further study. Indeed, the current findings linked to ‘GNAI2,P2RY12’ should be viewed with some caution as G coupled protein receptor interaction with G protein typically occurs inside the cell (Rosenbaum et al., 2009).

Beyond the three pairs mentioned above, our analysis also indicated other positive correlations. For example, the molecules associated with the ligand-receptor pair ‘VTN,CD47’ which emerged only in comparison between AD7 and T2D have been previously linked with neurodegeneration (Gheibihayat et al., 2021; Ruzha et al., 2022). The top 10 correlated pairs were subjected to preventive action inference, a procedure that utilized aging-related changes. It should be borne in mind that any assumption inferring that all age-associated changes are bad is not absolute and the direction of change needs to be taken into account. Among all actions inferred, the LRP1 on astrocyte stood out from all datasets.

### Combing with previous studies, LRP1 is a high potential target and needs further investigation

LRP1 is a multifunctional endocytic receptor involved in a broad spectrum of biological processes, including lipid metabolism, protein clearance and also cell signalling. LRP1 is known to interact with over 100 ligands (Mantuano et al., 2022). In the current CCC inference results, several ligands exhibited overlapping changes with LRP1, including PSAP, C1QB, APP, and APOE. Among these, AB is particularly important, as previous studies have demonstrated that LRP1 facilitates AB clearance in brain neurons via endocytosis, contributing to its role in AD disease pathophysiology (Kanekiyo & Bu, 2014). Molecular docking simulations using PyDock further suggested that APP displayed the strongest predicted binding affinity to LRP1 among the examined ligands. However, this result must be interpreted with caution, as the simulations were conducted using fragmented structural data rather than full-length protein interactions, which may not fully capture *in vivo* binding dynamics. Necessitating experimental validation using techniques such as cryo-electron microscopy (cryo-EM), cross-linking mass spectrometry, or mutagenesis studies would be beneficial. Actual protein level validation should also be considered such as FACS, ELISA, and Western blotting. Further research is required to establish the functional relevance of these binding interactions in disease progression and therapeutic targeting.

Previous studies have long recognized LRP1’s role in AB metabolism, both in its membrane-bound and soluble (sLRP1) forms. Indeed, soluble LRP1 has been shown to bind approximately 70% of circulating AB in plasma, functioning as a key component of AB clearance via molecular drainage mechanisms (Shibata et al., 2000). It is known that LRP1 protein levels can be upregulated by resveratrol treatment, despite no observed changes at the gene expression level, suggesting that post-transcriptional mechanisms may regulate LRP1 protein abundance (Santos et al., 2016). Moreover, studies examining AD patient plasma samples have revealed that sLRP1 levels are significantly reduced, and its ability to bind AB is compromised, thus potentially contributing to impaired AB clearance and neurodegeneration (Shibata et al., 2000). Although LRP1 is widely regarded as an AB clearance receptor, it has been found that its interaction with APP and other ligands may have bidirectional effects on AB homeostasis (Storck & Pietrzik, 2017). This suggests that LRP1’s role in neurodegeneration is more complex than initially believed, as it may act as both a neuroprotective and pathogenic factor depending on cellular context and disease stage (Storck & Pietrzik, 2017). The ability of LRP1 to modulate APP processing and AB production raises critical questions regarding how its function is regulated across different cell types and whether targeting LRP1 in therapeutic strategies could yield unintended effects. Resveratrol has been identified as a potentially therapeutic natural compound aiming age-related diseases by many studies (Gu et al., 2021; Jhanji et al., 2022; Rao et al., 2024; Yuan et al., 2024; Zhou et al., 2021). The study used bulk RNA data from normal murine brain, cross-referring with resveratrol-treated brain bulk RNA sequence data, only LRP1 gene emerged from both AD7 and AD6. Thus, LRP1 could potentially be the key modulator for age-related brain changes and also AD and T2D where it could be a potential target for disease prevention and treatment.

## Summary

The study has leveraged scRNA data and bioinformatic CCC inference approaches to investigate potential shared brain-linked neurodegenerative mechanisms across normal ageing and two age-related diseases, AD and T2D. These findings suggest that the APP-LRP1 interaction, in particular, plays a crucial role in CCC underlying age-related brain, particularly in microglia, endothelial cell, and astrocyte interactions. These results also highlight LRP1 as a potential therapeutic target that warrants further investigation, especially given its well-established role in AB clearance, neurodegeneration and neurovascular function. LRP1 remains an important but complex factor which is associated with brain neurodegeneration via its interactions with APP, Aβ, and other ligands (Shibata et al., 2000). The current study has provided foundation data upon which to build further mechanistic investigation, to fully understand the role of LRP1 in the brain, especially in relation to maintaining NVU homeostasis and neuronal function. Given its involvement in both neuroprotective and potentially pathogenic processes, future research would need to carefully delineate context-specific LRP1 functions with the possibilities of identifying novel pathways and targeted therapeutic development that maximise patient benefits while minimizing unintended disruptions to normal physiological processes.

