## Supplementary material for "Cell-cell communication analysis demonstrates early-stage common pathways linking ageing, Alzheimer’s disease, and Type 2 diabetes-related brain dysfunction": sup_figure

Sup fig1

1

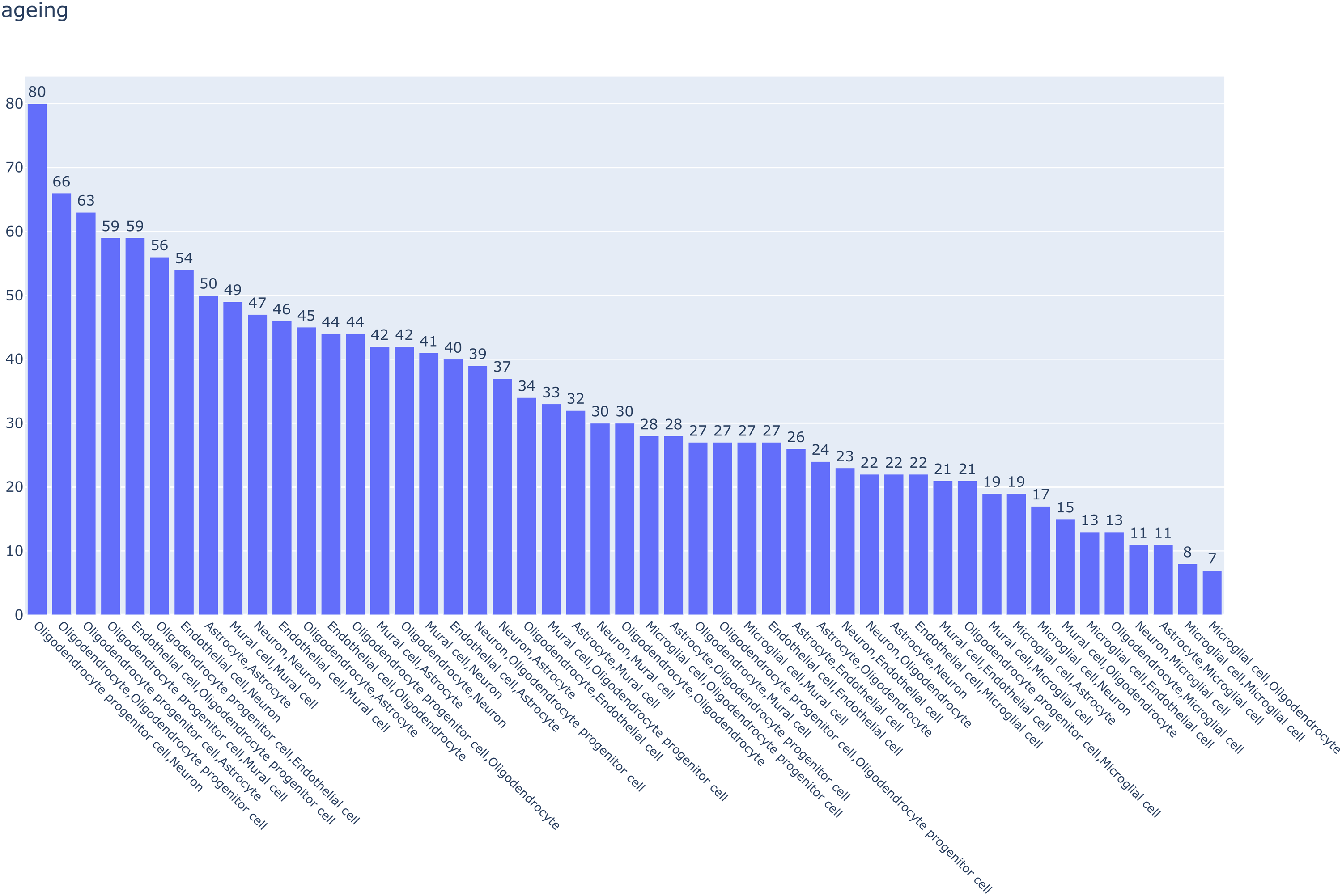

2
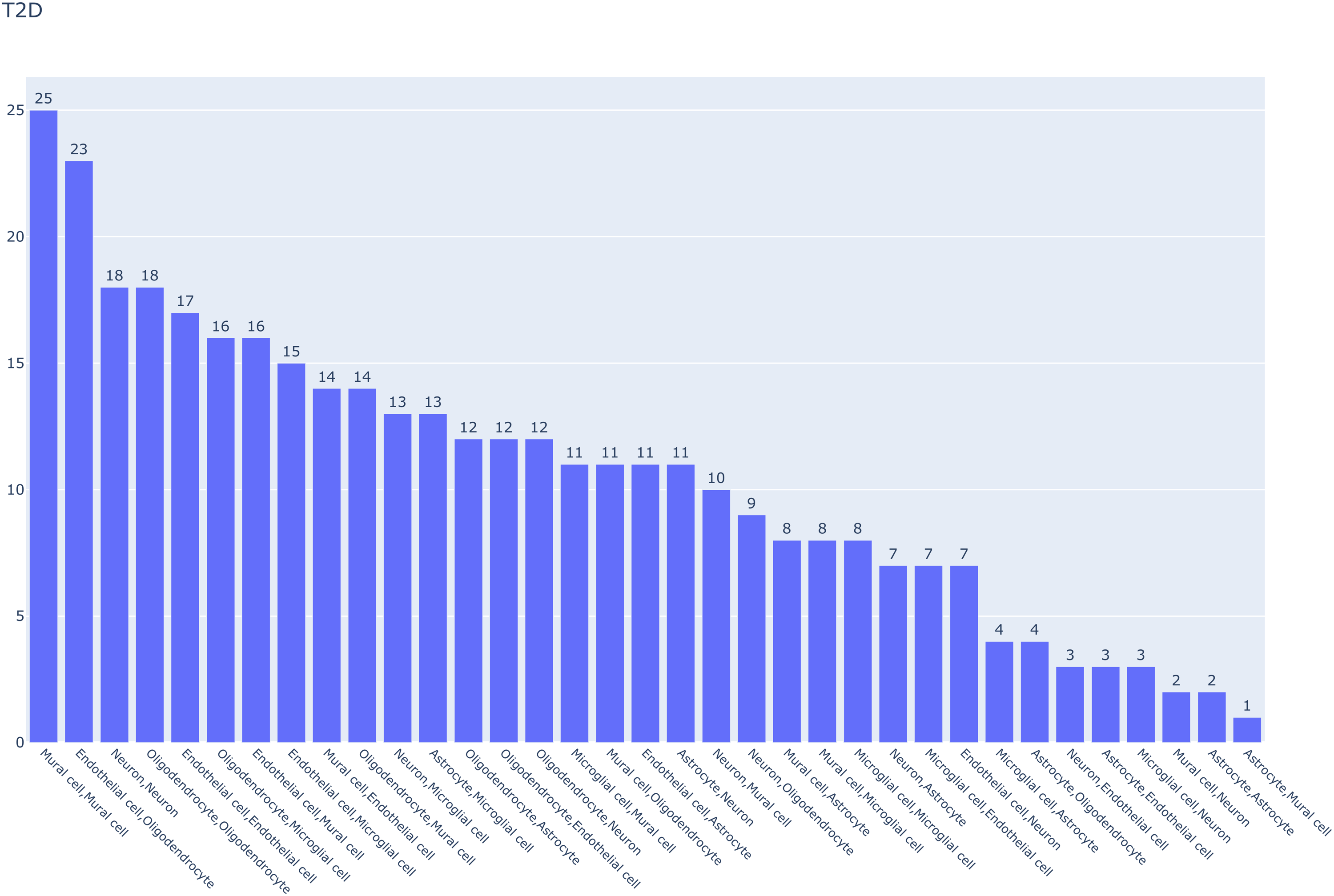
 3
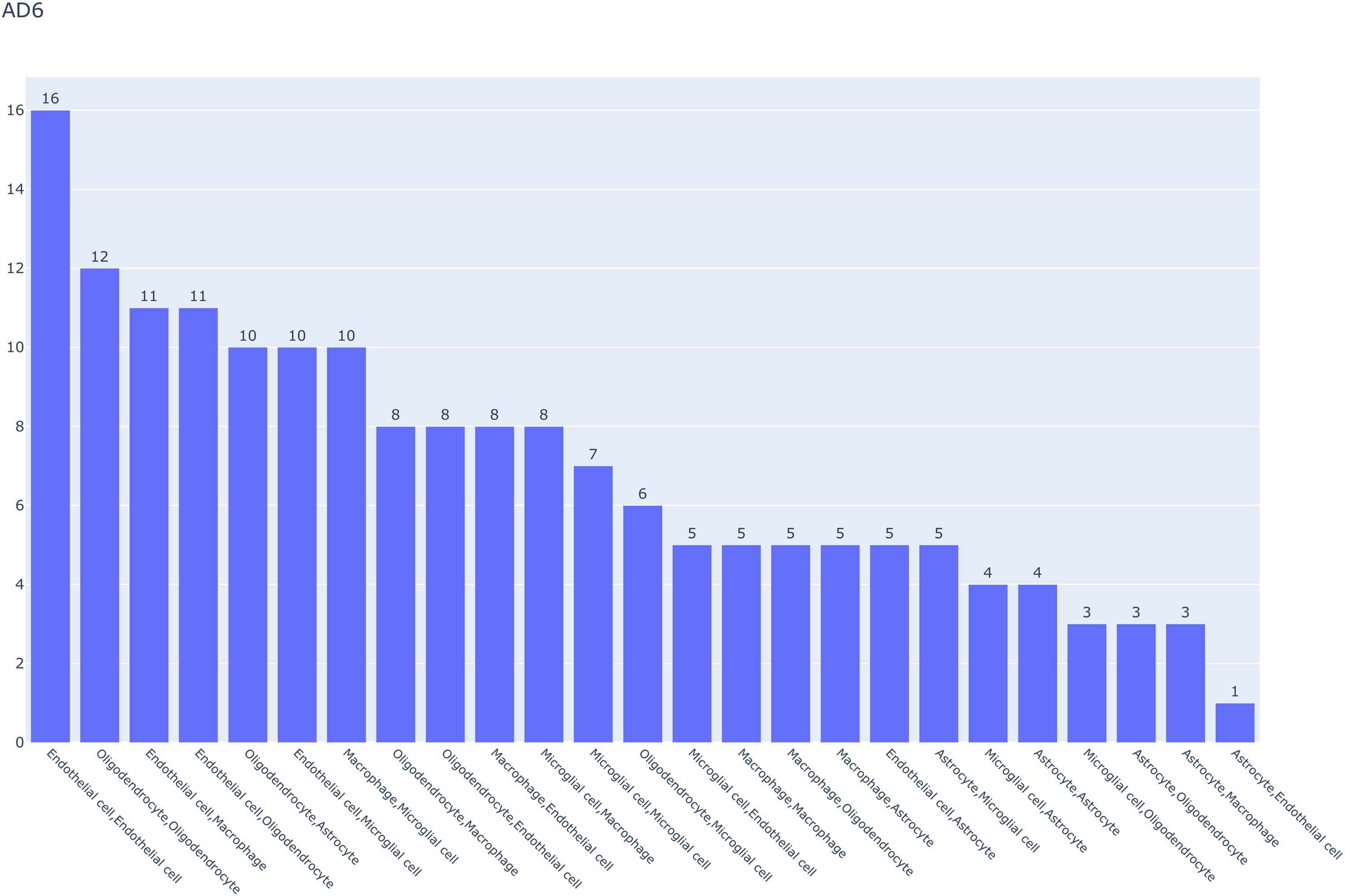
4

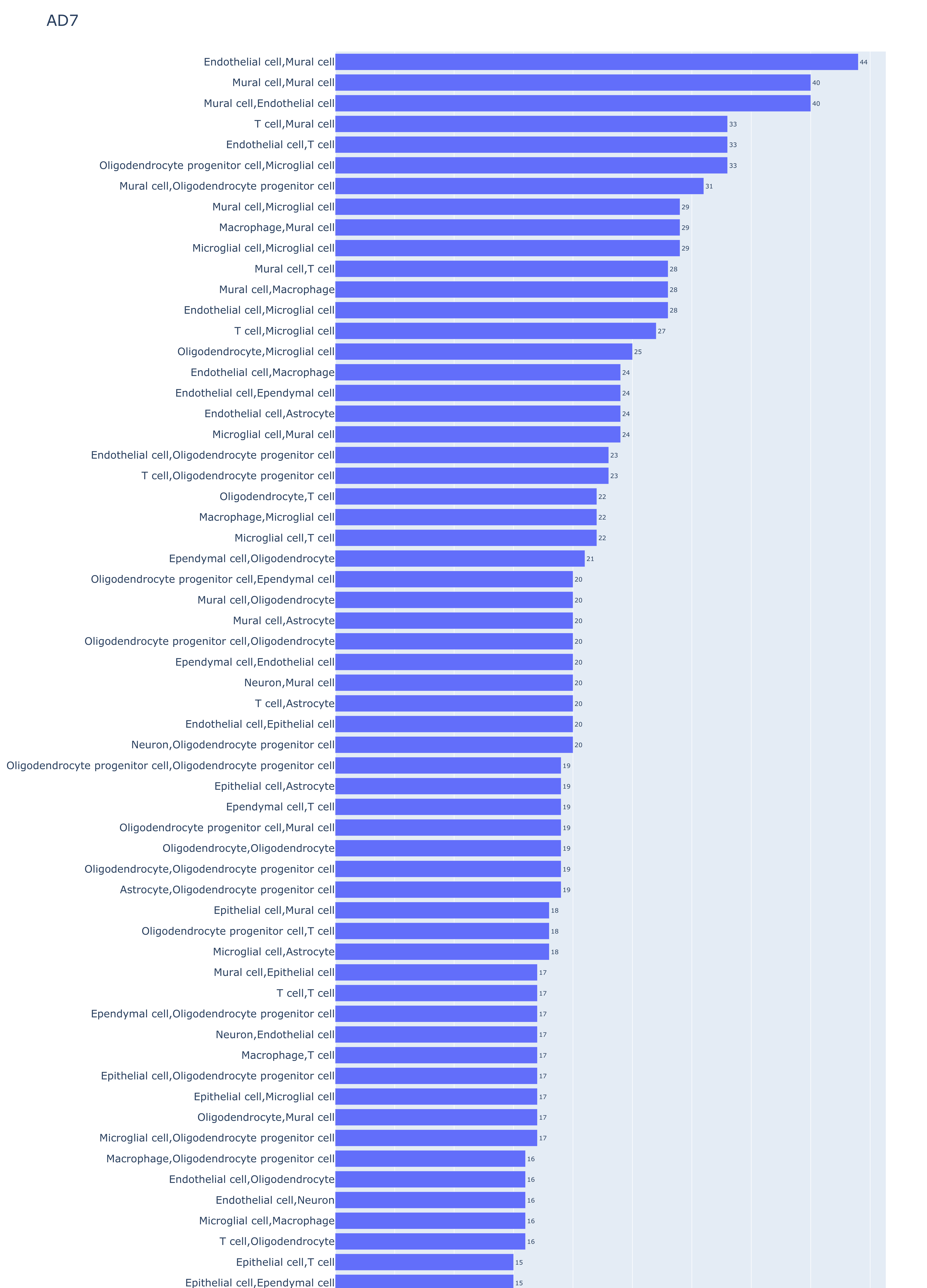

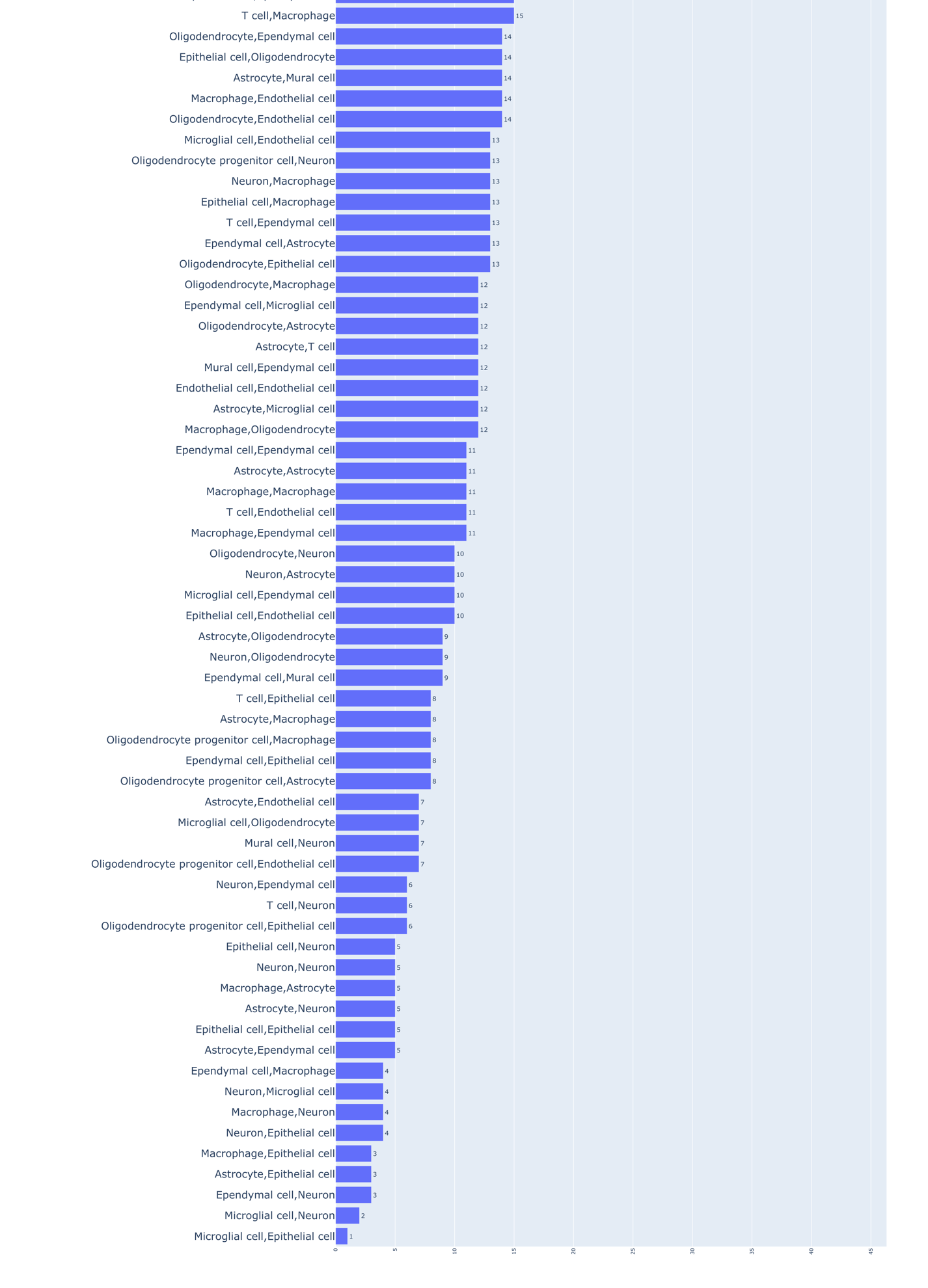

Sup figure1 the types of communication in each dataset ranked by the total number of changes regardless of change category. The bar indicated the magnitude of change for each type of communication.

Sup fig2

A

1
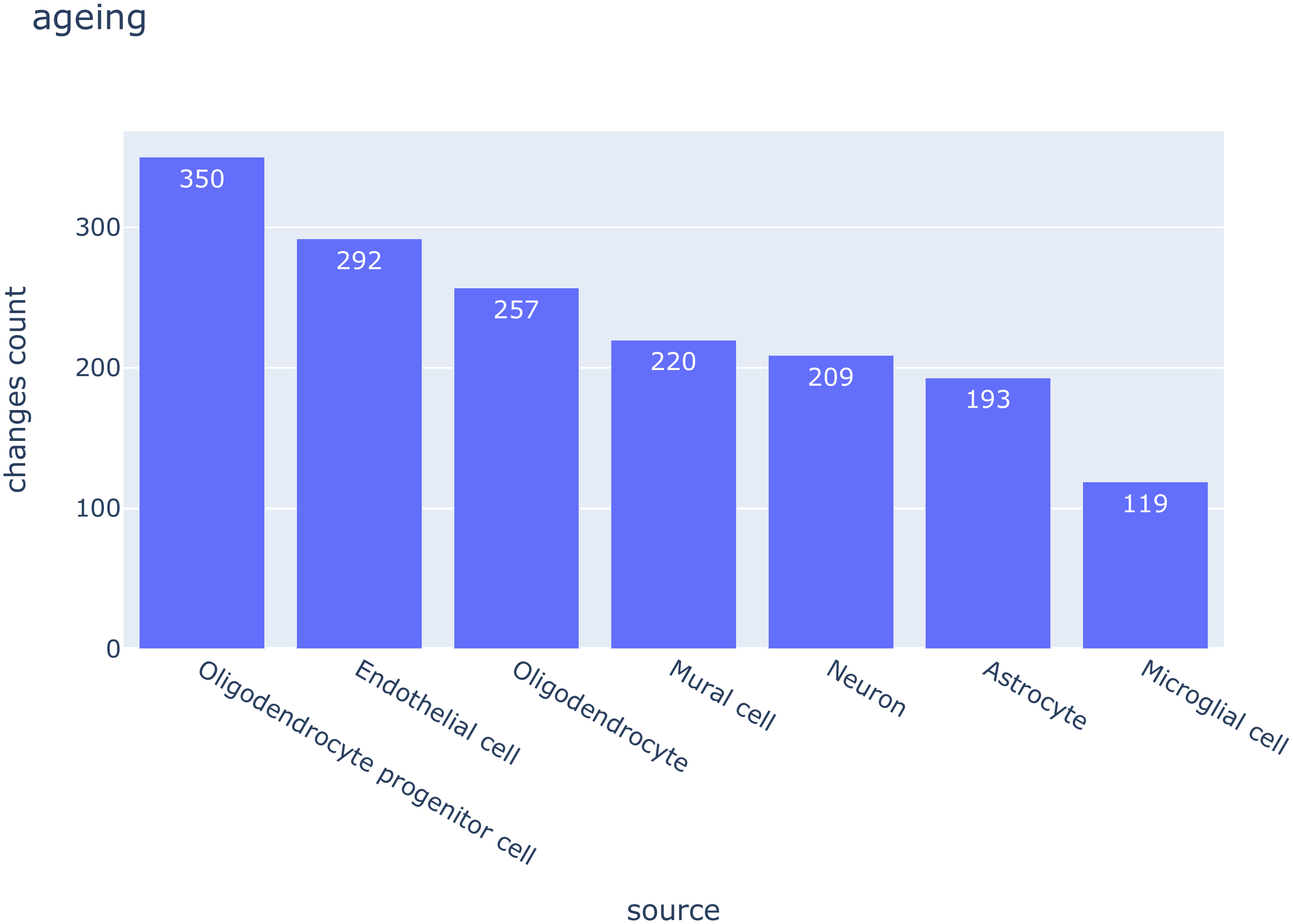
2
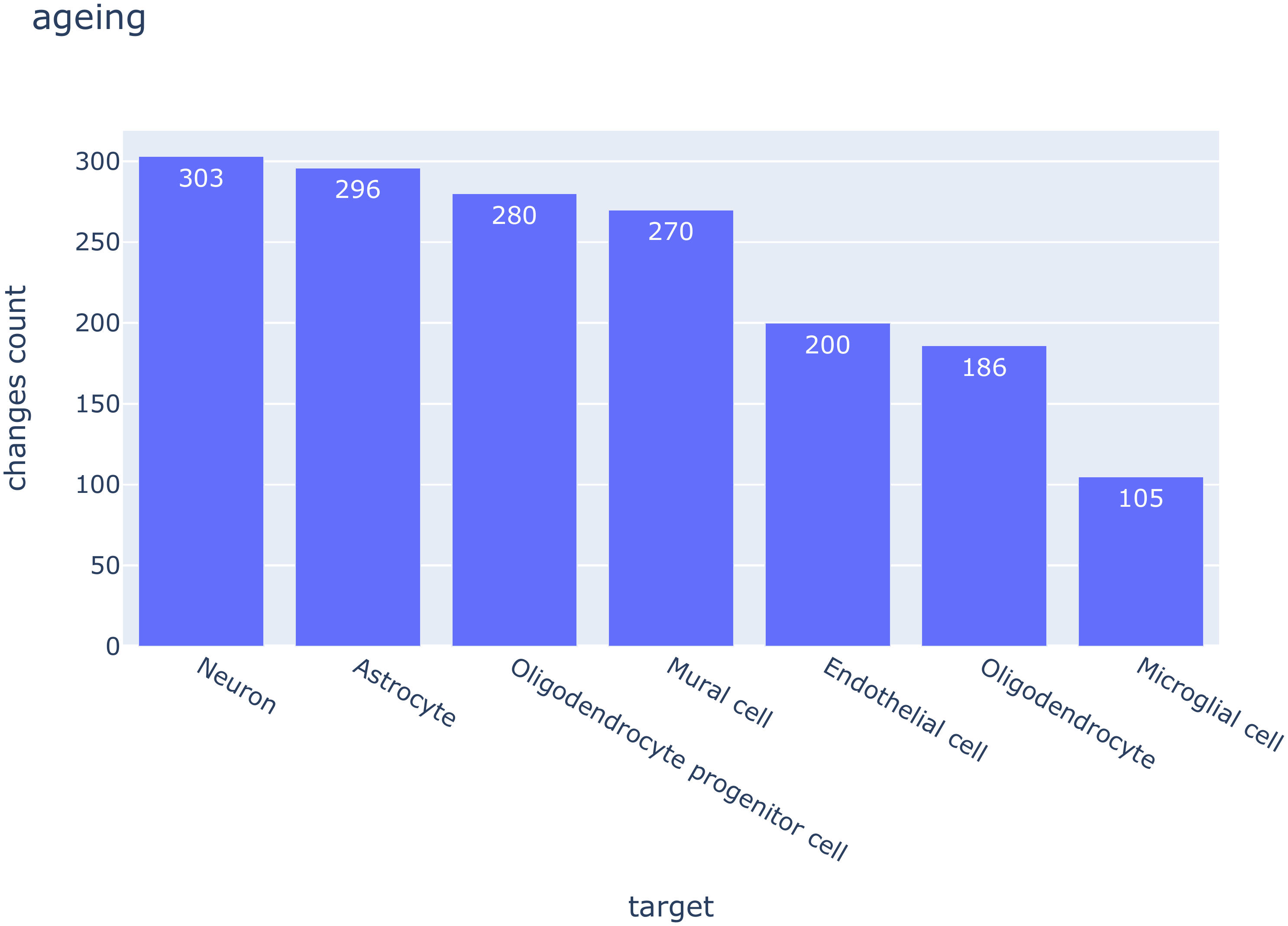
3

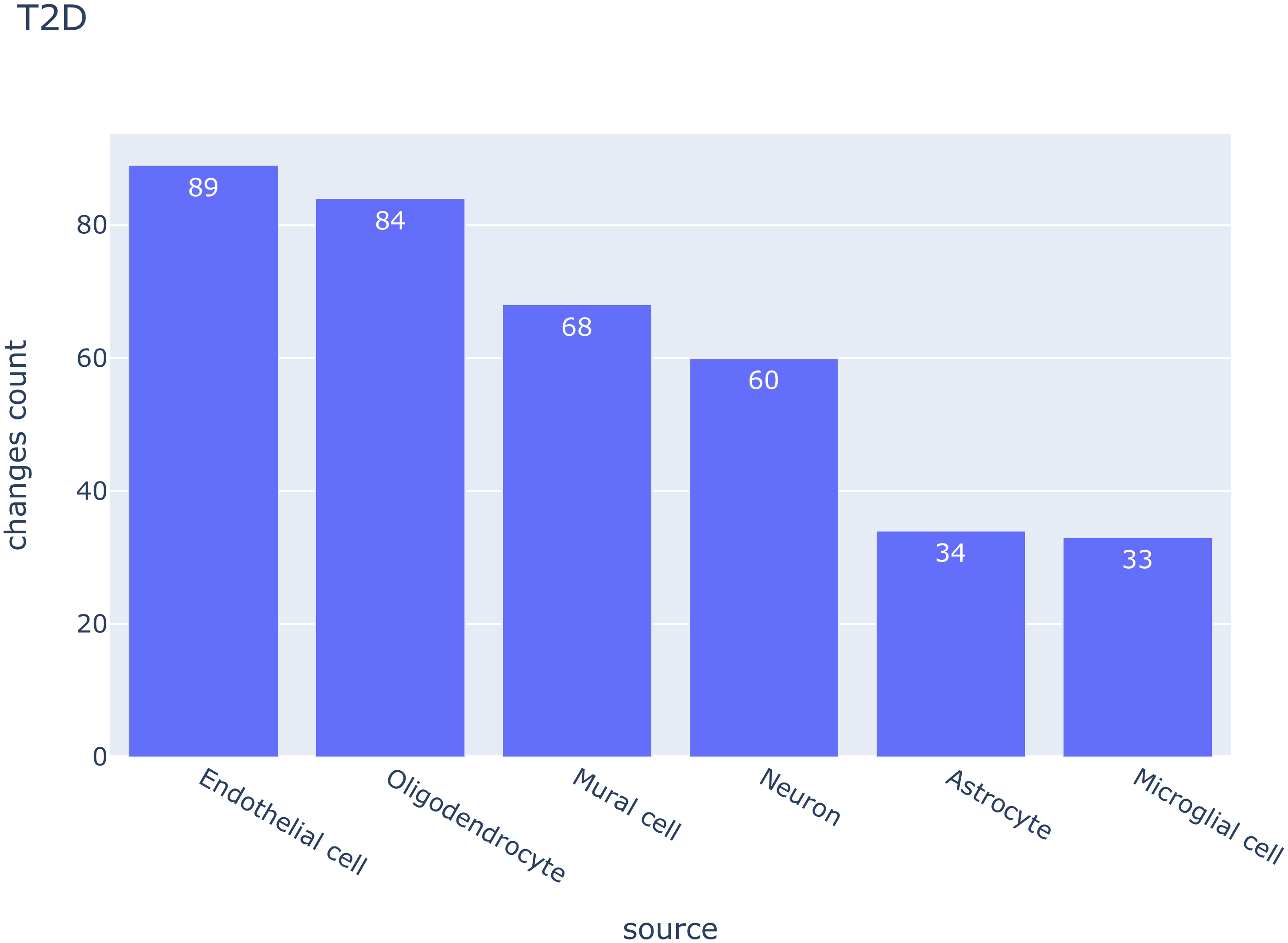
4

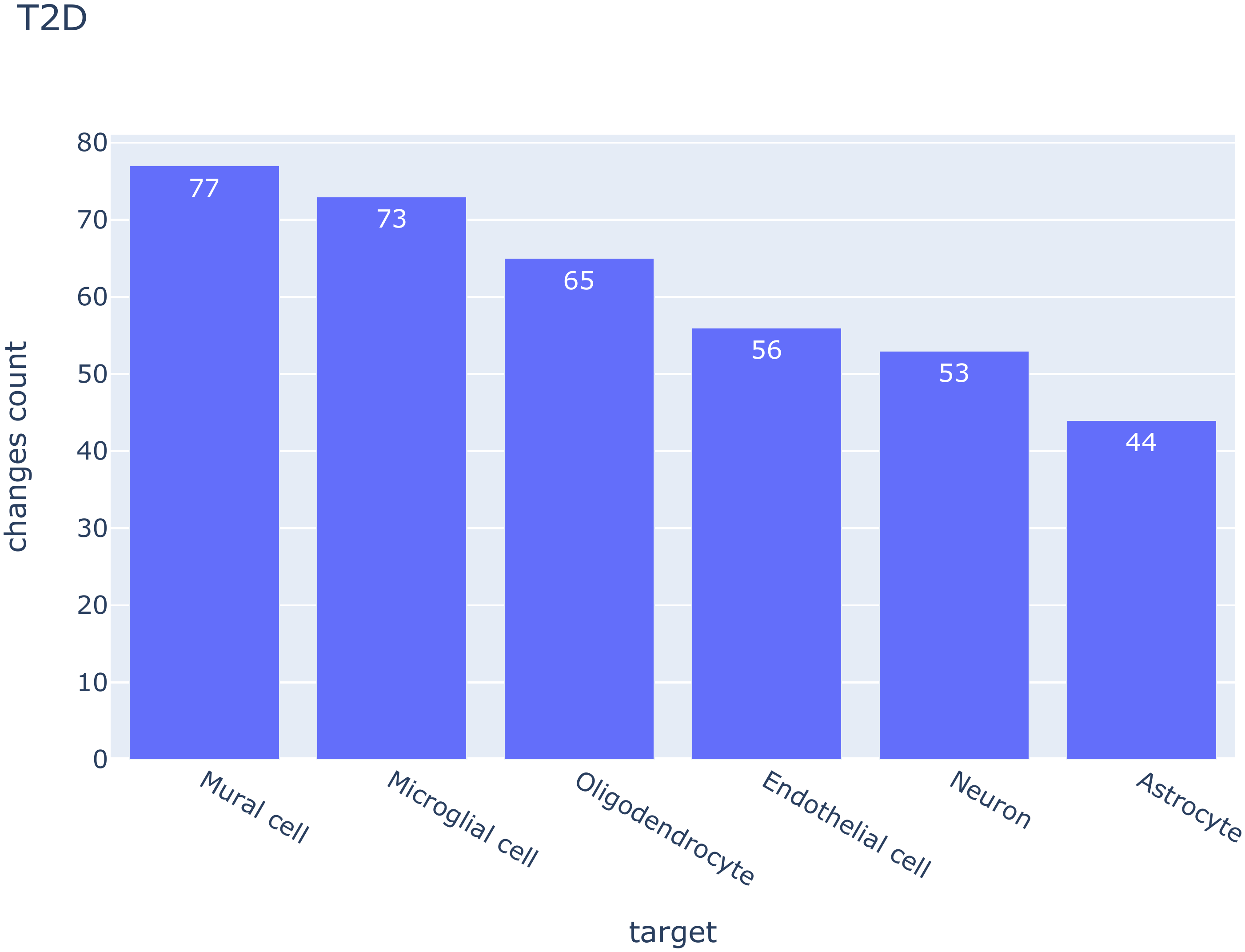

5
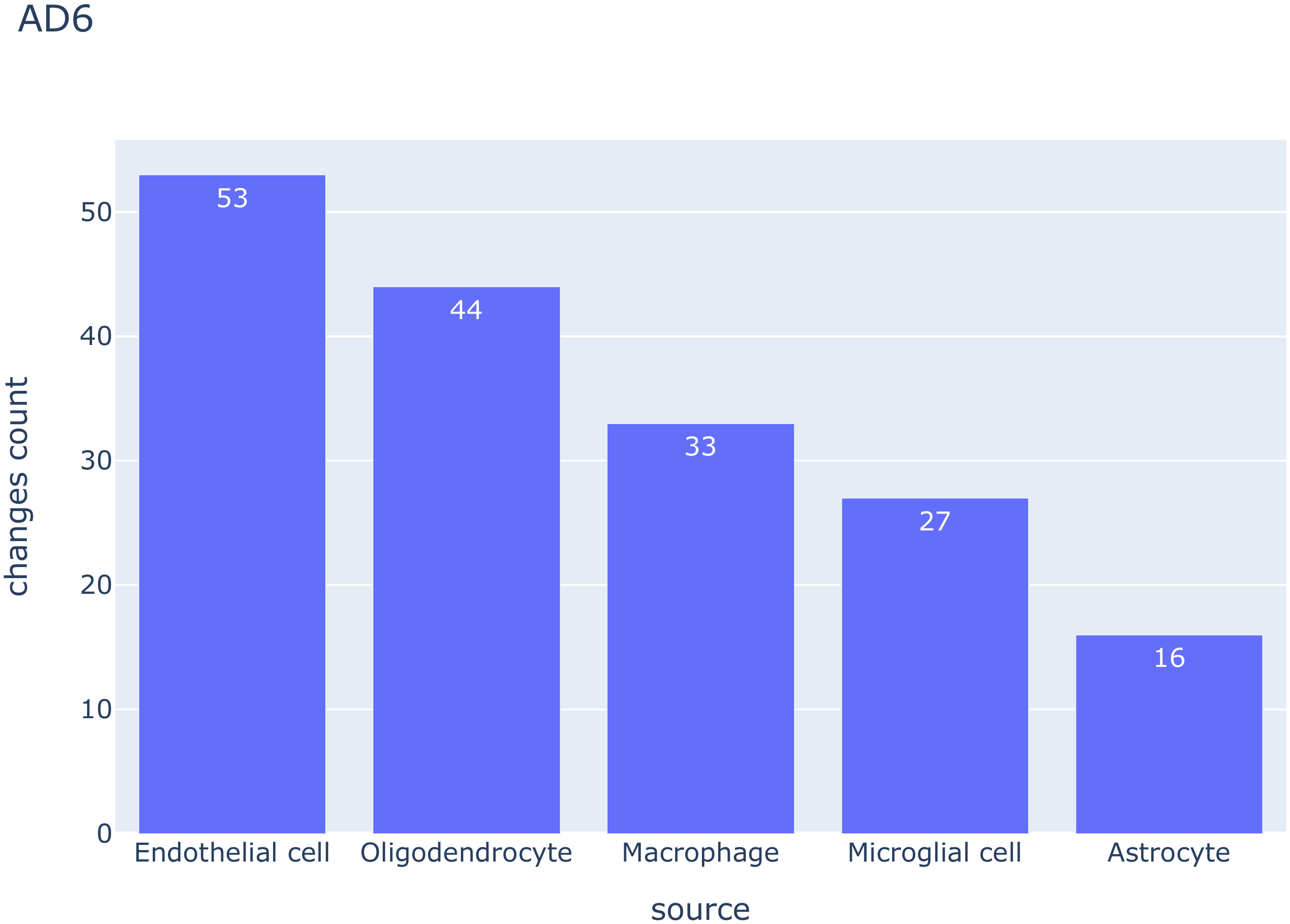
6
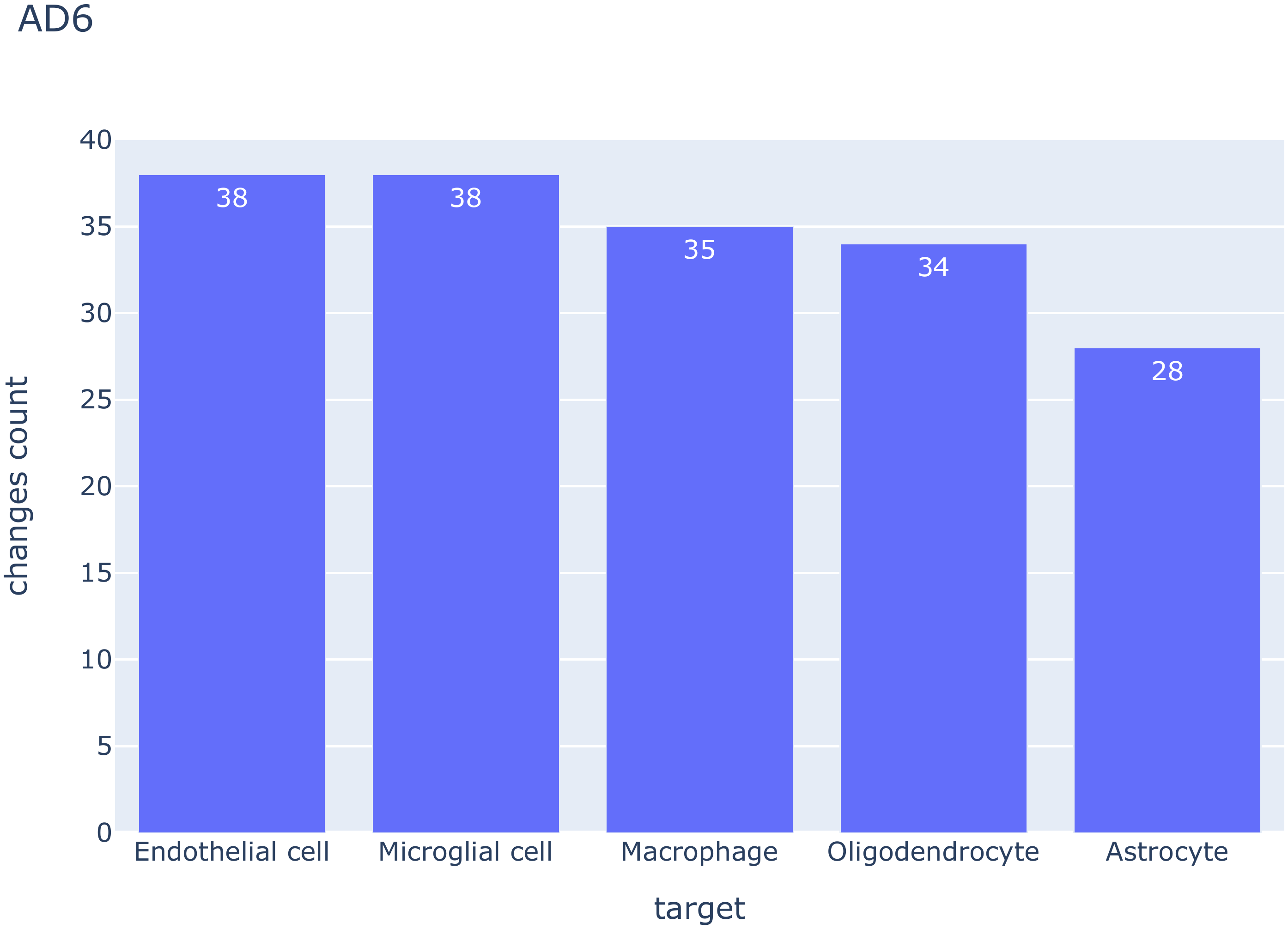
7
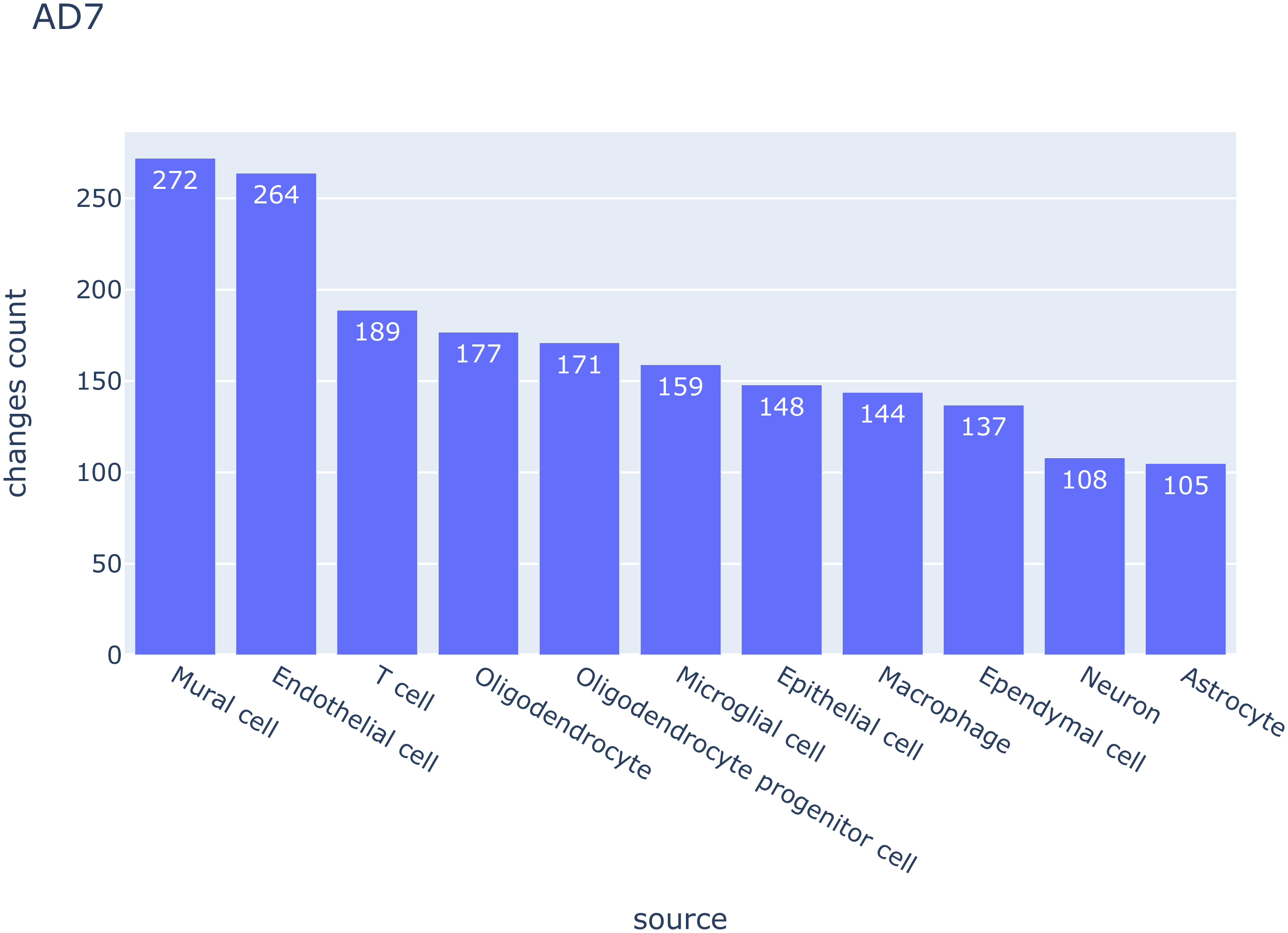
8
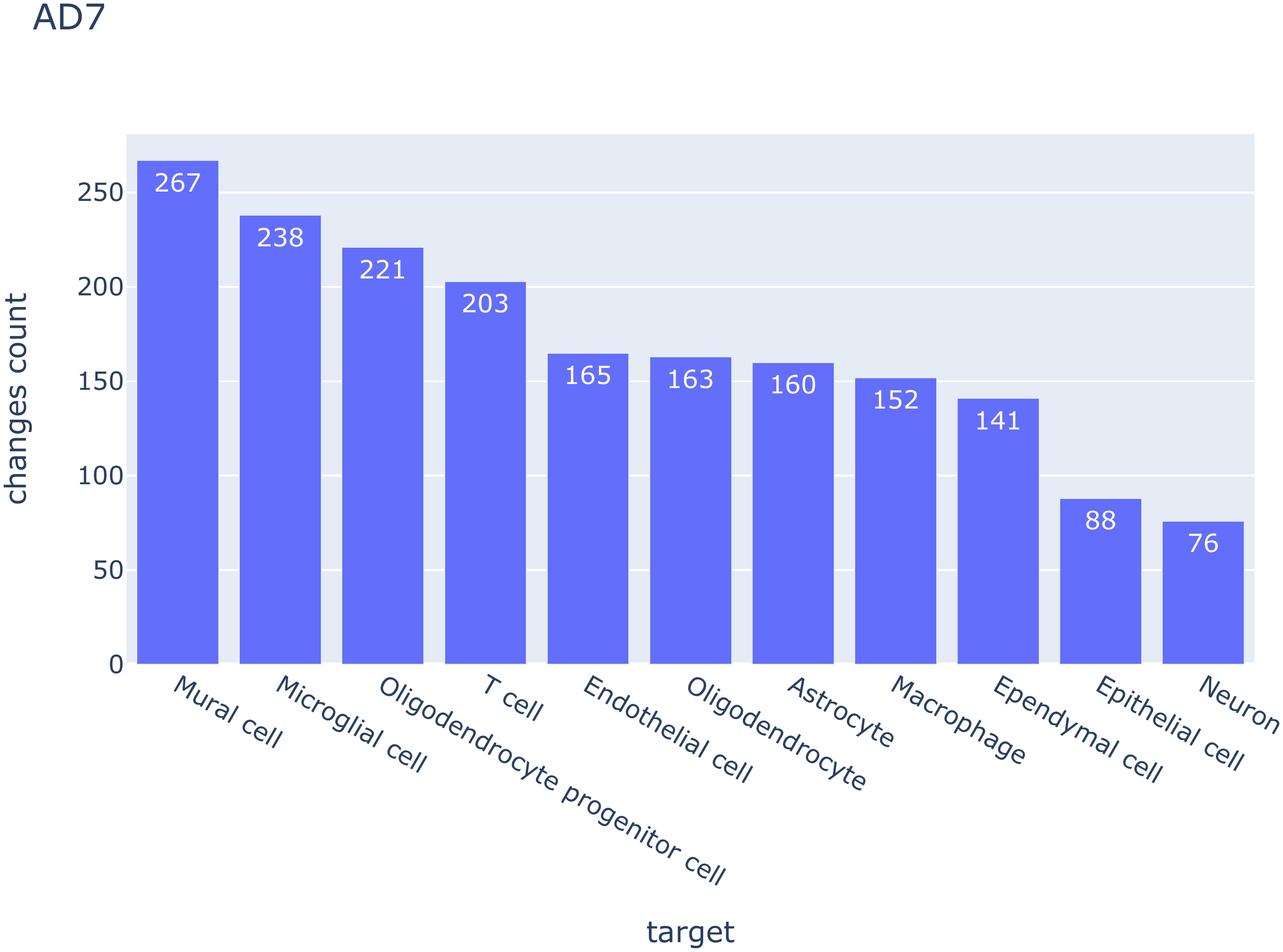

Sup figure 2 (A) cell types ranked by total number of changes as source cell types and target cell type respectively in each dataset. Source cell meant the cell type that gives the ligand while the target cell received signal by a certain receptor.
